# IL-1β turnover by TRIP12 and AREL1 ubiquitin ligases and UBE2L3 limits inflammation

**DOI:** 10.1101/2022.09.14.507790

**Authors:** Vishwas Mishra, Anna Crespo-Puig, Callum McCarthy, Tereza Masonou, Izabela Glegola-Madejska, Alice Dejoux, Gabriella Dow, Matthew J. G. Eldridge, Luciano H. Marinelli, Meihan Meng, Shijie Wang, Daniel J. Bennison, Avinash R. Shenoy

## Abstract

The cytokine interleukin-1β (IL-1β) has pivotal roles in antimicrobial immunity, but also incites inflammatory pathology. Bioactive IL-1β is released following proteolytic maturation of the pro-IL-1β precursor by caspase-1 inflammasomes. UBE2L3/UBCH7, a conserved ubiquitin conjugating enzyme, promotes pro-IL-1β ubiquitylation and proteasomal disposal. However, UBE2L3 actions *in vivo* and ubiquitin ligases involved in this process are unknown. Here we report that deletion of *Ube2l3* in mice markedly reduces pro-IL-1β turnover in macrophages, leading to excessive mature IL-1β production, neutrophilic inflammation and disease symptoms following inflammasome activation. A family-wide siRNA screen identified two ubiquitin ligases, TRIP12 and AREL1, which we show add K27-, K29- and K33- poly-ubiquitin chains on lysine residues in the ‘pro’ domain and destabilise pro-IL-1β. Mutation of ubiquitylation sites increased pro-IL-1β stability, but did not affect proteolysis by caspase-1. The extent of mature IL-1β production is therefore determined by precursor abundance, and UBE2L3, TRIP12 and AREL1 limit inflammation by shrinking the cellular pool of pro-IL-1β. Our study has uncovered fundamental processes governing IL-1β homeostasis and provided molecular insights that could be exploited to mitigate its adverse actions in disease.

## INTRODUCTION

Interleukin-1β (IL-1β) has been at the forefront of cytokine biology for over 50 years, yet the name is a misnomer because IL-1β does much more than ‘communicate between leukocytes’ [1]. Apart from enhancing the functions of myeloid and lymphoid cells during infection, IL-1β promotes systemic inflammation by activating acute-phase protein production by hepatocytes, prostaglandin release, enhanced coagulation and neutrophil adherence on endothelial cells, and anti-viral activity in keratinocytes [1]. In addition, IL-1β is an endogenous pyrogen that induces fever at doses as low as 1 ng.kg^-1^ body weight in humans. Deregulated IL-1β production drives the pathology of autoinflammatory diseases such as gout, rheumatoid arthritis, diabetes, cardiac and neurodegenerative disease, and hereditary fever syndromes [1]. Given these pivotal roles of IL-1β, we need a better understanding of processes governing its production and new approaches to neutralise its detrimental actions.

IL-1β production is tightly controlled transcriptionally and post-translationally [1, 2]. In myeloid cells (e.g., macrophages, monocytes), which are the dominant source of IL-1β, *Il1b* mRNA is induced in an NF-κB—dependent manner following exposure to microbial molecules (e.g., ligands of Toll-like receptors) or cytokines (e.g., TNF) [1]. *Il1b* mRNA codes for the biologically inert pro-IL-1β precursor protein (270 aa). Pro-IL-1β undergoes proteolysis by caspase-1, originally called interleukin-1 converting enzyme, which removes the ‘pro’ domain (aa 1-116) and triggers the release of the mature, receptor-binding cytokine (aa 117-270; hereon we refer to the precursor as pro-IL-1β and mature form as IL-1β). Despite the wealth of information on bioactive IL-1β, regulation of the pro-IL-1β precursor prior to its proteolysis by caspase-1 remains poorly understood.

Caspase-1 is activated within inflammasomes, which are multimolecular signalling scaffolds [2]. Inflammasomes detect cytosolic microbial ligands (e.g., cytosolic LPS [3]) or ‘sterile’ danger signals (e.g., gout-associated uric acid crystals [4]), among other triggers. LPS, a component of Gram-negative bacteria, induces systemic inflammation through its potent ability to induce *Il1b* transcripts and pro-IL-1β conversion via caspase-4/11, NLRP3 (NOD and leucine rich repeat containing protein with a pyrin domain 3) and caspase-1. Inflammasome components such as IL-1β, NLRP3, caspase-1, and caspase-11 are thus among the major drivers of septic-shock like disease induced by LPS in mice [5]. Similarly, genetic loss of *Casp1*, *Nlrp3*, or *Casp11* abolishes IL-1β production and neutrophilic inflammatory symptoms induced by sterile signals, such as alum, cholesterol or uric acid crystals [1, 2, 5].

How cells manage pro-IL-1β abundance or dispose of the precursor cytokine is not completely understood. We previously discovered that the ubiquitin conjugating enzyme UBE2L3 (also called UBCH7) promotes pro-IL-1β ubiquitylation and proteasomal degradation in macrophages [6]. Ubiquitylation involves the covalent conjugation of ubiquitin to proteins, followed by further additions on one or more lysine residues of ubiquitin to form poly-ubiquitin chains [7, 8]. This requires ubiquitin activating E1 enzymes (2 in mice and humans), ubiquitin conjugating E2 enzymes (∼40 in mice and humans) and ubiquitin E3 ligases (∼700 in mice and humans). UBE2L3 is among the most abundant E2 enzymes in cells and only partners with E3 ligases of the Homologous to E6 C-terminus (HECT) and Really Interesting New Gene (RING) between RING (RBR) subfamilies [9, 10]. These ubiquitin ligases transfer ubiquitin from the active site cysteine of UBE2L3 to their own catalytic cysteine residue before ubiquitylating substrates [7, 8]. UBE2L3 does not partner with the RING subfamily which directly transfer ubiquitin from E2 enzymes to substrates. This suggests that UBE2L3-dependent pro-IL-1β ubiquitylation likely involves HECT or RBR ligases, however, the precise proteins involved are not known.

UBE2L3 is an indirect target of caspase-1 and its abundance reduces upon inflammasome activation [6]. Silencing UBE2L3 expression enhances pro-IL-1β abundance in macrophages *in vitro* and its overexpression increases pro-IL-1β turnover, which indicates a key role of UBE2L3 in pro-IL-1β homeostasis [6]. In humans, polymorphisms in *UBE2L3* are linked to inflammatory conditions, including arthritis and inflammatory bowel disease, demonstrating a link between UBE2L3 and deregulated inflammation [10]. Whether UBE2L3 abundance is affected by inflammasomes *in vivo* and the consequence of *Ube2l3-*deletion in whole animals has not been tested. Here we generated and characterised conditional tissue-specific deletion of *Ube2l3* in mice (*Ube2l3^ΔMac^*), which resulted in elevated pro-IL-1β abundance and secretion of IL-1β following inflammasome activation. Unbiased RNA interference (RNAi) screening identified two HECT-type E3 ligases, TRIP12 and AREL1, in promoting pro-IL-1β ubiquitylation and proteasomal turnover. These findings provide insights on the regulation of a potent proinflammatory cytokine and E3 ligases that could be exploited in designing novel therapeutics in the future.

## RESULTS

### Inflammasome activation depletes macrophage UBE2L3 in vivo

Inflammasome activation in human and mouse macrophages *in vitro* results in UBE2L3 targeting for degradation, which facilitates higher IL-1β production [6]. We examined this *in vivo* by treating mice with LPS or LPS and ATP to activate the NLRP3 inflammasome as described previously [11] and assessing UBE2L3 abundance in peritoneal macrophages. Both treatments led to IL-1β production, but as expected, the administration of LPS and ATP led to more robust inflammasome activation and higher IL-1β secretion as compared to LPS alone (*Figure S1A*). Flow cytometry of peritoneal macrophages revealed reduced UBE2L3 staining (*Figure S1B*) and geometric mean fluorescence intensity (*Figure S1C*) in macrophages from mice given LPS or LPS and ATP as compared to PBS as negative control. Consistent with this, immunoblots of macrophages from three representative mice showed lower UBE2L3 abundance (*Figure S1D*). These experiments verified that UBE2L3 abundance reduces in macrophages upon inflammasome activation *in vivo*.

UBE2L3 levels correlate inversely with IL-1β production *in vitro* as it promotes the proteasomal disposal of pro-IL-1β [6]. Similarly, an inverse correlation was found between UBE2L3 levels in peritoneal macrophages and IL-1β in peritoneal lavage fluid (Pearson’s correlation coefficient r = -0.7, n = 15, *P* value: 0.0039; *Figure S1E*). Taken together, these results indicate that UBE2L3 is depleted by inflammasomes and could act as a negative regulator of IL-1β production *in vivo*.

### Generation of conditional Ube2l3-deletion in macrophages (Ube2l3^ΔMac^)

To investigate the functions of UBE2L3 *in vivo,* we generated a tissue-specific conditional deletion strain because *Ube2l3* is an essential gene in mice [12]. Mice with a ‘floxed’ *Ube2l3* exon 1 (*Ube2l3^fx/fx^* mice) were crossed with Csf1r-cre/Esr1 mice that express the Cre recombinase-oestrogen receptor fusion protein whose activity is induced by 4-hydroxytamoxifen (hTam) in Csf1r+ve cells (i.e., macrophages and monocytes, hereafter called *Ube2l3^ΔMac^* mice; *Figure 1A*) [13]. hTam treatment induced recombination at the *Ube2l3* locus as verified by genomic DNA PCR in *Ube2l3^ΔMac^* primary bone marrow-derived macrophages (BMDMs) (*Figure S2A*) and dramatically reduced UBE2L3 protein levels as assessed by immunoblotting (*Figure 1B*). These experiments established the conditional knockout system to examine the role of UBE2L3.

**Figure 1.**
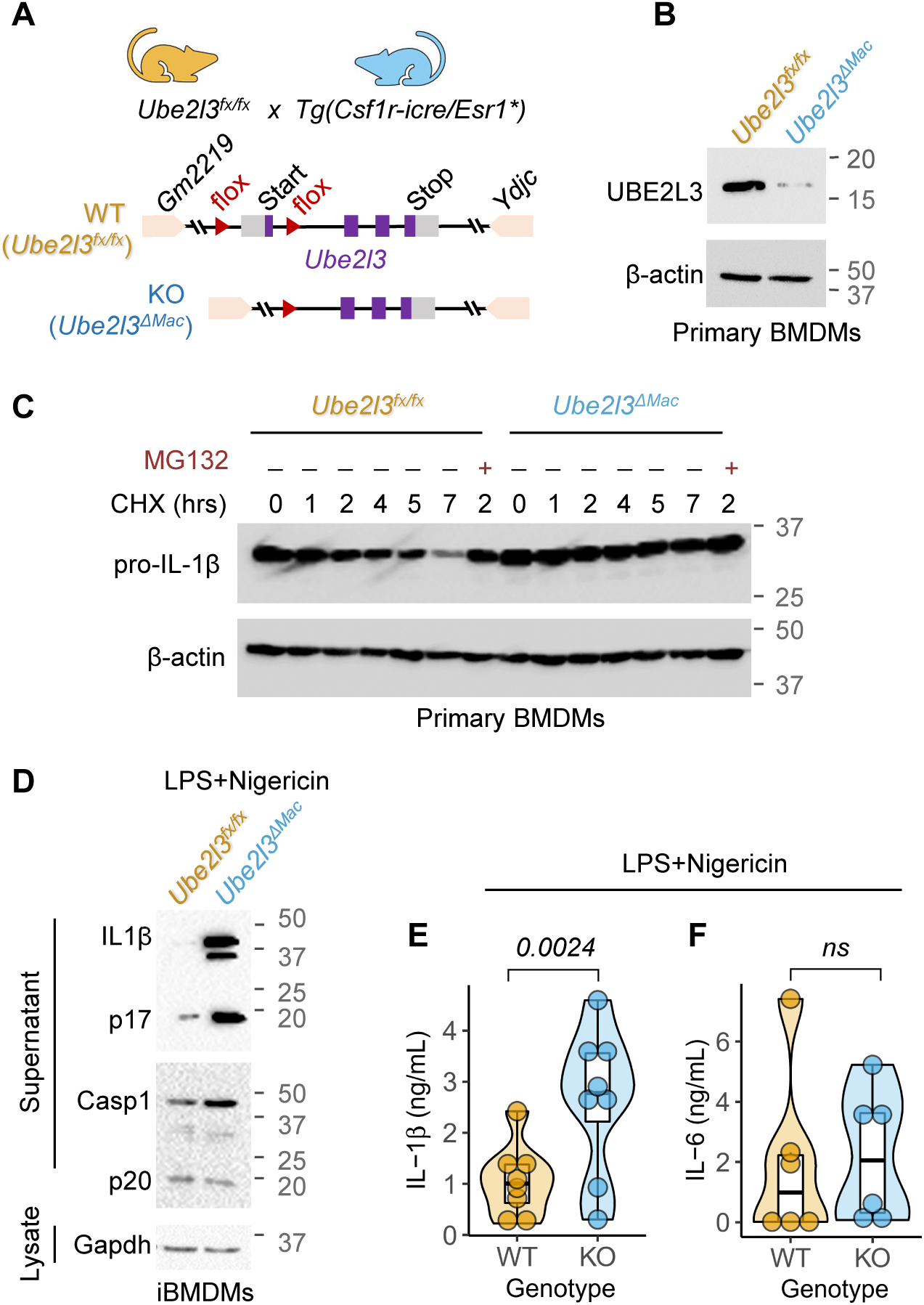
*Ube2l3* deletion increases pro-IL-1β stability in macrophages. **(A)** Schematic depiction of the ategy to generate conditional, macrophage-specific deletion of *Ube2l3*. Exon 1 of *Ube2l3* was ‘floxed’, ding to its deletion by Cre recombinase whose expression is controlled by the Csf1r (also called CD115) omoter and activity is induced by oral administration of tamoxifen *in vivo* or 4-hydroxytamoxifen *in vitro* in acrophages. **(B)** Representative immunoblots for UBE2L3 and β-actin from lysates of primary BMDMs of the dicated genotypes treated with 4-hydroxytamoxifen (2 μM, 48 h). **(C)** Representative immunoblots of pro-1β and β-actin in cell lysates from cycloheximide (CHX)-chase experiments carried out with primary DMs of the indicated genotypes given 4-hydroxytamoxifen (2 μM, 48 h). Cells were treated with LPS (250 .mL^-1^) for 14 h followed by CHX (10 μg.mL^-1^) for indicated times. MG132 (10 μM) was added for the last 3 h in e indicated samples. **(D)** Representative immunoblots of cell lysates and supernatants of iBMDMs of the dicated genotypes given 4-hydroxytamoxifen (2 μM, 48 h) followed by LPS (250 ng.mL^-1^, 3 h) and nigericin 0 μM, 1 h) to activate inflammasomes. **(E-F)** Quantification of IL-1β **(E)** and IL-6 **(F)** by ELISA of supernatants m iBMDMs of the indicated genotypes treated as in **D**. WT = *Ube2l3^fx/fx^*; KO = *Ube2l3*^!1^*^Mac^*. Immunoblots are presentative of 2 independent experiments in **B-C** and 3 independent experiments in **D**. Each dot in **E-F** presents a biologically independent repeat. Data distribution is depicted with violin, box (25th to 75th rcentile, line at median), and whiskers (± 1.5 x IQR); data are pooled from 2-3 independently repeated periments with and age- and sex-matched mice. Two-tailed *P* values for indicated comparisons from mixed ects ANOVA; ns = not significant (*P* > 0.05).

To examine how *Ube2l3* loss-of-function affected pro-IL-1β turnover, we performed cycloheximide (CHX)-chase experiments, in which inhibition of protein translation with CHX reveals stability of proteins over time (‘chase’). CHX-chase on LPS-treated primary BMDMs from *Ube2l3^fx/fx^* and *Ube2l3^ΔMac^* given hTam revealed that pro-IL-1β protein turnover was markedly slower in *Ube2l3^Δmac^* macrophages as compared to wild-type macrophages (*Figure 1C*). Proteasome inhibition with MG132 blocked pro-IL-1β clearance (*Figure 1C*). Together, this indicated UBE2L3 is required for proteasomal pro-IL-1β turnover.

In line with high pro-IL-1β levels, *Ube2l3^ΔMac^* immortalised BMDMs (iBMDMs) treated with LPS and nigericin to activate the NLRP3 inflammasome secreted prominently higher mature IL-1β as measured by ELISA and immunoblots (*Figure 1D, E*). Caspase-1 activation and IL-6 secretion were similar in both genotypes (*Figure 1D, 1F*), indicative of normal NF-κB signalling, inflammasome priming and activation. We therefore conclude that deletion of *Ube2l3* reduces pro-IL-1β protein turnover and does not affect inflammasome activation. Importantly, these results from knockout mice unequivocally establish a crucial role for UBE2L3 in reducing pro-IL-1β abundance.

### Higher circulating IL-1β and disease severity in Ube2l3^Δmac^ mice after inflammasome activation

We first assessed pro-IL-1β turnover in peritoneal macrophages isolated from *Ube2l3^ΔMac^* mice given tamoxifen to induce knockout *in vivo*. Efficacy of tamoxifen-induced silencing of UBE2L3 expression *in vivo* in *Ube2l3^ΔMac^* mice was verified by flow cytometry and immunoblots of peritoneal macrophages (*Figure 2A; S2B*). Indeed, pro-IL-1β turnover was slower in *Ube2l3^ΔMac^* than *Ube2l3^fx/fx^* peritoneal macrophages in CHX-chase experiments (*Figure 2B*). Importantly, LPS-induced *Il1b* mRNA measured by qRT-PCR was similar in both genotypes (*Figure S2C*), which indicates that UBE2L3 controls pro-IL-1β abundance post-translationally.

**Figure 2.**
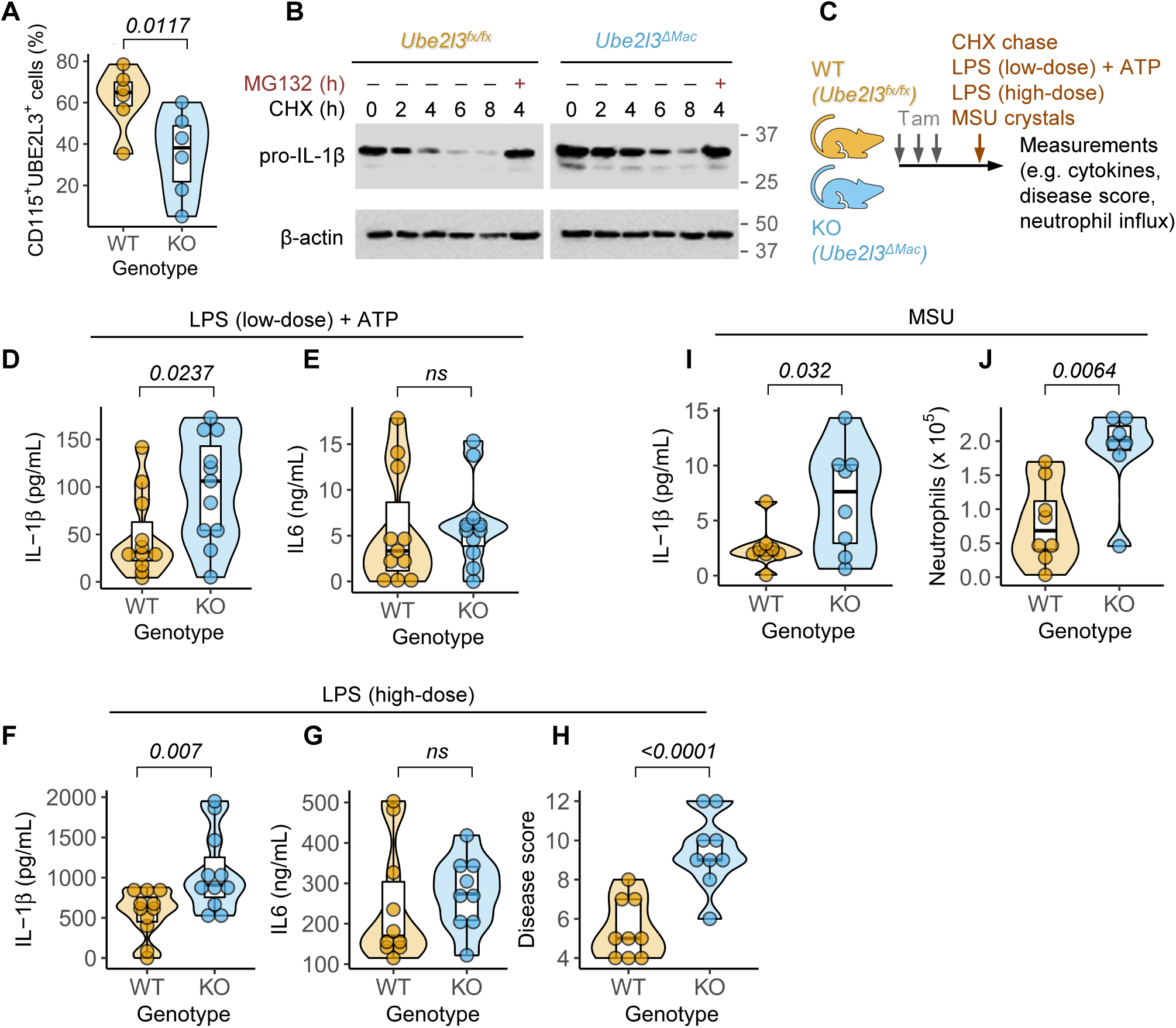
Elevated IL-1β in *Ube2l3****^Δ^****^Mac^* mice after inflammasome activation. **(A)** Flow cytometry showing percentage of Csf1r/CD115 and UBE2L3 +ve peritoneal macrophages isolated from mice of the indicated genotypes given tamoxifen orally on 3 consecutive days. WT = *Ube2l3^fx/fx^*; KO = *Ube2l3^ΔMac^*. **(B)** Representative immunoblots of pro-IL-1β and β-actin in cell lysates from cycloheximide (CHX)-chase experiments carried out on peritoneal macrophages isolated from the indicated mice given tamoxifen. Cells were treated with LPS (250 ng.mL^-1^) for 14 h followed by CHX (10 μg.mL^-1^) for indicated times. MG132 (10 μM) was added for the last 3 h in the indicated samples. Images from the same exposure of the same immunoblots are shown with intervening lanes removed. **(C)** Schematic depiction of inflammatory models tested in *Ube2l3^fx/fx^* (WT) and *Ube2l3^ΔMac^* (KO) mice as indicated. Mice were given tamoxifen (Tam; 80 mg.kg^-1^) via oral gavage on 3 consecutive days, and the indicated inflammatory stimuli intraperitoneally on day 5. Low dose LPS+ATP = 5 μg LPS per mouse for 3 h followed by ATP (50 μmol) for 10 min. High dose LPS = 30 mg.kg^-1^ LPS for 3 h. MSU crystals = monosodium uric acid crystals (1 mg per mouse) for 6 h. See Methods for details. **(D-E)** Quantification of IL-1β **(D)** and IL-6 **(E)** by ELISA of peritoneal lavage from mice of the indicated genotypes following low dose LPS model. WT = *Ube2l3^fx/fx^*; KO = *Ube2l3^ΔMac^*. **(F-H)** Quantification of IL-1β **(F)** and IL-6 **(G)** by ELISA of serum, and disease scores **(H)**, of mice of the indicated genotypes following high-dose LPS treatment. WT = *Ube2l3^fx/fx^*; KO = *Ube2l3^ΔMac^*. **(I-J)** Quantification of IL-1β **(I)** by ELISA and flow cytometry-based counts of neutrophils **(J)** in the peritoneal lavage of mice injected with MSU crystals. WT = *Ube2l3^fx/fx^*; KO = *Ube2l3^ΔMac^*. Immunoblots in **B** represent experiments carried out at least two times independently from two mice each of the indicated genotypes. Each dot in **D-J** represents a mouse. Data distribution is depicted with violin, box (25th to 75th percentile, line at median), and whiskers (± 1.5 x IQR); data are pooled from 2-4 independently repeated experiments. Two-tailed P values for indicated comparisons from mixed effects ANOVA. ns –not significant (*P* > 0.05).

Based on these results, we hypothesised that inflammasome activation will result in higher circulating IL-1β and inflammation in *Ube2l3^ΔMac^* mice. To test this, *Ube2l3^fx/fx^* and *Ube2l3^ΔMac^* mice given tamoxifen were tested in three models of inflammasome— dependent inflammation to quantify IL-1β production and inflammatory parameters (*Figure 2C*). In the low-dose LPS model as above, IL-1β in the peritoneal lavage was 2-fold higher in tamoxifen-treated *Ube2l3*-deficient mice as compared to *Ube2l3-*sufficient control animals (*Figure 2D*). Secretion of the inflammasome-independent IL-6 cytokine was similar in both groups of mice (*Figure 2E*), indicating a specific effect of *Ube2l3-*deletion on IL-1β production.

As LPS injection serves as a model of severe Gram-negative bacterial infection-associated sepsis we also tested a high-dose LPS model that results in septic shock-like disease. In this model as well *Ube2l3^ΔMac^* mice had 2-fold higher serum IL-1β compared to *Ube2l3^fx/fx^* mice (*Figure 2F*); IL-6 levels remained unchanged (*Figure 2G*). In agreement with high IL-1β levels, endotoxic shock-like disease activity scores (see Methods) were twice as higher in mice lacking *Ube2l3* (*Figure 2H*). These experiments together established a critical role for macrophage UBE2L3 in antagonising IL-1β production and dampening inflammatory disease symptoms *in vivo*.

We next tested whether UBE2L3 is also involved in a model of low-grade sterile inflammation. Sterile monosodium uric acid crystals (MSU) elicit NLRP3-dependent activation of caspase-1 and IL-1β release, which then stimulates neutrophil influx into the peritoneum [4]. As compared to wild-type mice injected with MSU, *Ube2l3^ΔMac^* mice had 3-fold higher IL-1β (*Figure 2I*) and 2-fold greater neutrophil influx in the peritoneum (*Figure 2J*), which further underscore the role of UBE2L3 in suppressing IL-1β production *in vivo*.

Taken together, these experiments establish that conditional deletion of *Ube2l3* in macrophages/monocytes is sufficient to trigger elevated IL-1β, local and systemic inflammation upon inflammasome activation, revealing its remarkably powerful role in limiting IL-1β—driven inflammation. Alongside earlier experiments in knockout macrophages, we conclude that UBE2L3 promotes pro-IL-1β turnover, and in its absence the increased accumulation of pro-IL-1β becomes available for caspase-1—mediated maturation.

### RNAi screen for E3 ligases that promote pro-IL-1β turnover

What is the mechanism underlying UBE2L3-dependent pro-IL-1β turnover? As UBE2L3 is a ubiquitin-conjugating enzyme, we sought to identify the ubiquitin E3 ligase(s) that it co-opts for pro-IL-1β ubiquitylation. We previously showed that the abundance of pro-IL-1β protein in macrophages reduces between 9-24 h following LPS-treatment and proteasomal inhibitors MG132 or epoxomicin block pro-IL-1β turnover [6]. To narrow down the potential E3 ligases that could be involved in this process, we performed similar experiments in the presence of broad-specificity inhibitors of subclasses of E3 ligases. Firstly, BAY-11-7082, which can inhibit UBE2L3 and RBR E3 ligases within the linear ubiquitin assembly complex (LUBAC) [14], among other targets, increased pro-IL-1β abundance as compared to vehicle (DMSO)-treated cells (*Figure S3A*). Members of the Cullin-RING superfamily of E3 ligases are activated by the attachment of NEDD8 to Cullin, a step that can be blocked by MLN4924 which inhibits the NEDD8-activating E1 enzyme [15]. However, treatment with MLN4924 did not affect pro-IL-1β abundance (*Figure S3A*), ruling out the involvement of Cullin-RING E3 ligases. Heclin, an inhibitor of the HECT-type E3 ligases NEDD4, SMURF2 and WWP1 [16], also did not affect pro-IL-1β abundance (*Figure S3A*). Based on these results we designed a family-wide siRNA screen against 43 HECT and RBR E3 ligases to identify the protein(s) involved.

Pro-IL-1β expression is induced by NF-κB activation by TLR ligands (e.g., LPS) or proinflammatory cytokines (e.g., TNF) through signalling steps that involve ubiquitylation/deubiquitylation (e.g., LUBAC, TRAFs, A20) [1, 2]. For the RNAi screen, we wanted to specifically measure pro-IL-1β protein abundance over time and therefore engineered a doxycycline-inducible pro-IL-1β expression system that is independent of NF-κB signalling. Doxycycline dose-dependent expression of pro-IL-1β^strep^ (pro-IL-1β with two C-terminal StrepTag II tags) was detected in iBMDMs stably transduced with the pLTREK2P-pro-IL-1β^strep^ lentiviral plasmid (*Figure S3B-C*). In these cells, doxycycline-induced pro-IL-1β^strep^ expression was abrogated by siRNA-mediated silencing of *rtta3*, the transcription factor encoded by the pLTREK2P plasmid (*Figure S3B*), which confirmed the specificity of doxycycline-controlled expression. We also verified that the mechanism and kinetics of pro-IL-1β^strep^ turnover were similar to that of endogenous pro-IL-1β in several independent ways. Firstly, the abundance of LPS-induced endogenous pro-IL-1β and doxycycline-induced pro-IL-1β^strep^ declined similarly over time (*Figure S3D*). The same inhibitors that blocked endogenous pro-IL-1β turnover as above also blocked pro-IL-1β^strep^ turnover: i.e., only inhibitors of the proteasome (MG132) or UBE2L3 (BAY-11-7082) reduced pro-IL-1β^strep^ levels (*Figure 3A*). Most importantly, RNAi-mediated silencing of UBE2L3 expression increased the abundance of doxycycline-induced pro-IL-1β^strep^ (*Figure 3B*). These results together endorsed the doxycycline-inducible pro-IL-1β^strep^ as a faithful reporter for our RNAi screen.

**Figure 3.**
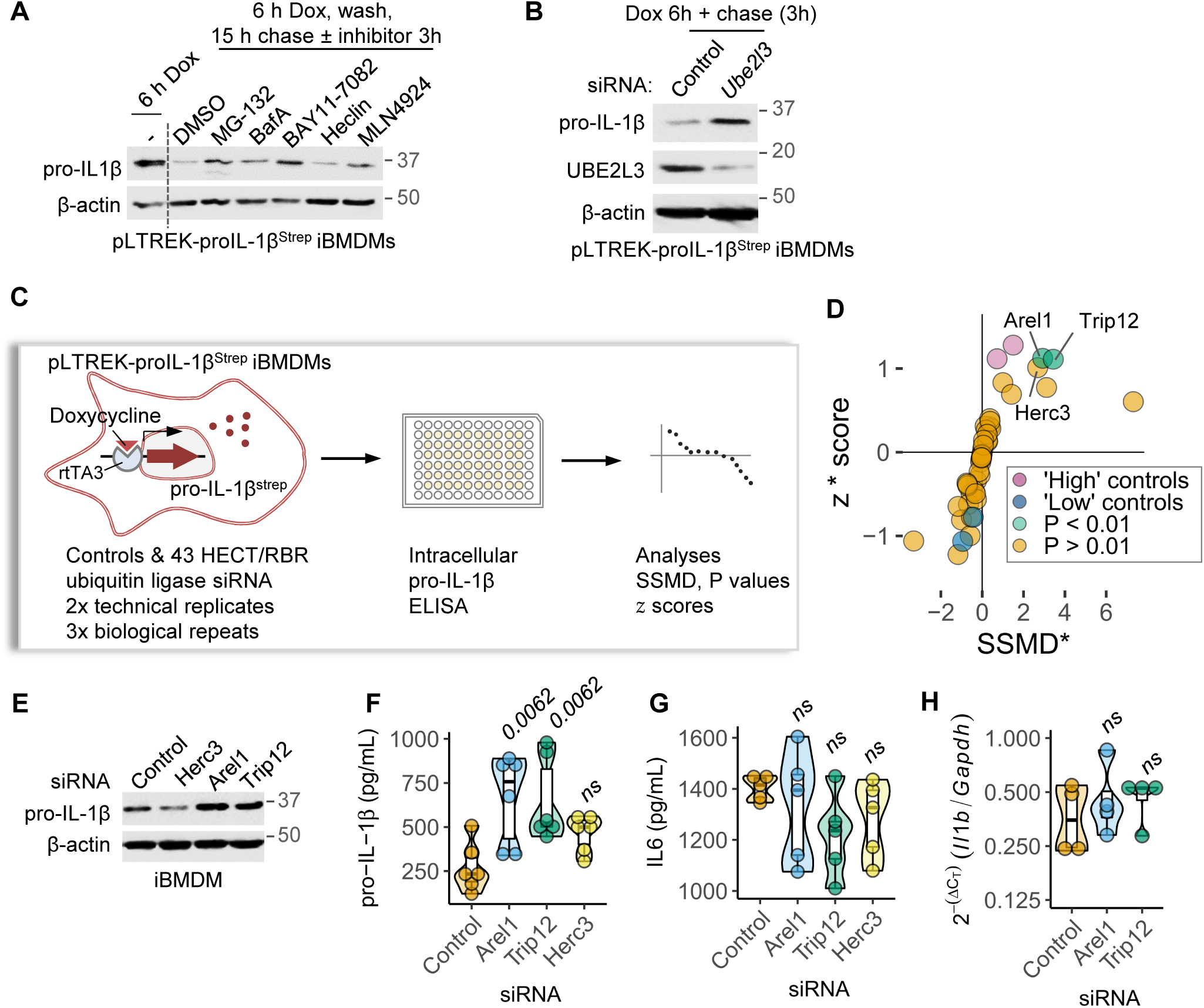
siRNA screen identifies the E3 ligases TRIP12 and AREL1 in promoting pro-IL-1β turnover. **(A)** resentative immunoblots of cell lysates from iBMDMs stably transduced with pLTREK-proIL-1βStrep plasmid wing the effect of the indicated inhibitors on the abundance of doxycycline (Dox)-induced pro-IL-1βStrep. s were treated with Dox for 6 h, washed, and incubated for 18 h (‘chased’) and the indicated inhibitors or the cle DMSO was added for the last 3 h. Cell lysate collected at 6 h (sample labelled (—)) served as a control. 32 (proteasome inhibitor) and BAY 11-7082 (LUBAC and UBE2L3 inhibitor) treatments increased pro-IL-trep abundance as compared to DMSO or other treatments. **(B)** Representative immunoblots showing the ct of silencing UBE2L3 expression on the abundance of doxycycline (Dox)-induced pro-IL-1βStrep. pLTREK-L-1βStrep iBMDMs were transfected with non-targeting control or UBE2L3 siRNA for 72 h, followed by Dox tment followed by a 3 h chase as indicated. **(C)** Schematic depiction of the siRNA screen against 43 HECT RBR E3 ligases in iBMDMs stably transduced with the pLTREK2-pro-IL-1β^Strep^ plasmid. The screen was ied out in 96-well plates and was independently repeated three times with technical duplicates within each at. Cells were transfected with siRNA for 72 h, treated with doxycycline, and intracellular pro-IL-1βStrep was ntified by ELISA. z* score, SSMD* and *P* values were calculated as described in Methods. **(D)** Plot showing the lt from siRNA screen described in **C**. Mean z* score and SSMD* from three independent repeats are plotted. trols are depicted in different colours as indicated in the legend. Trip12, Arel1, Herc3 and key controls are icted in colours and/or labelled. Symbols represent other genes and are coloured based on *P* values (see hods for further details). **(E-H)** Validation of hits from the siRNA screen described in **C-D** in iBMDMs sfected with non-targeting controls or siRNA against the indicated E3 ligases for 72 h followed by LPS (250 mL^-1^) treatment for 18 h. Representative immunoblots for endogenous pro-IL-1β **(E)** and quantification of acellular pro-IL-1β **(F)** and secreted IL-6 **(G)** by ELISA, and relative expression of *Il1b* transcript normalised to *dh* **(H)** are shown. Images in **A-B**, **E** represent experiments done at least three times. Each dot in **D** esents one gene or control conditions in the siRNA screen. In **F-H** each dot represents a biologically independent experiment. Data distribution is depicted with violin, box (25th to 75th percentile, line at median), and whiskers (± 1.5 x IQR). Two-tailed P values for comparisons with samples given Control siRNA in F-H from mixed effects ANOVA. ns – not significant (P > 0.05).

Family-wide siRNA screening was performed against 43 E3-ligases of the HECT and RBR family to determine their role in the turnover of pro-IL-1β^strep^ (*Figure 3C*; see Methods). Wells treated with MG-132 for last 3 h served as positive controls (’high’ pro-IL-1β^strep^ levels), and wells given siRNA against *rtta3* as negative controls (’low’ pro-IL-1β^strep^ levels; *Figure 3D, S3E*). Additional controls included wells untreated with doxycycline or without siRNA. Intracellular pro-IL-1β was quantified by ELISA on siRNA screens performed three times independently with technical replicates within each attempt (see Methods). Two hits stood out based on z* score, SSMD* and *P* value cut-offs ([17]; see Methods). These were two HECT-type E3 ligases: TRIP12 and AREL1 (*Figure 3D, S3E*).

### TRIP12 and AREL1 promote pro-IL-1β turnover

We next directly tested whether the E3 ligases identified in the screen contributed to the turnover of endogenous pro-IL-1β. In agreement with results from the screen, abundance of LPS-induced endogenous pro-IL-1β was reduced upon silencing TRIP12 or AREL1 as assessed by immunoblots and ELISA of intracellular pro-IL-1β (*Figure 3E-F, S3F*). Secretion of IL-6 and *Il1b* transcription remained unaffected by TRIP12 or AREL1 silencing (*Figure 3G-H*), showing that these proteins do not broadly affect NF-κB—dependent cytokine expression. Notably, silencing the next potential ‘hit’ based on SSMD* score, HERC3, had no effect on pro-IL-1β abundance or IL-6 production (*Figure 3E-G, S3F*), underscoring that the high stringency cut-off in the screen was appropriate.

We next asked whether silencing TRIP12 and AREL1 reduced pro-IL-1β turnover in CHX-chase assays. Indeed, silencing these E3 ligases increased pro-IL-1β stability (*Figure 4A, S3F*), which is a phenocopy of *Ube2l3-*deletion. We reasoned that elevated pro-IL-1β abundance upon silencing TRIP12 or AREL1 would lead to increased IL-1β release by inflammasomes. Indeed, macrophages given siRNA against TRIP12 or AREL1 released ∼2-fold more IL-1β following inflammasome activation with LPS and nigericin as compared to cells given non-targeting control siRNA (*Figure 4B*); as expected, the release of IL-6 remained unchanged (*Figure 4C*). Pyroptotic membrane damage measured by propidium iodide dye uptake assays was also unaffected (*Figure 4D*), indicating similar levels of NLRP3 inflammasome-driven pyroptosis. TRIP12 and AREL1 therefore do not affect inflammasome priming and activation. Overexpression of UBE2L3 in macrophages (iBMDM^YFP-UBE2L3^) increases pro-IL-1β ubiquitylation and turnover [6]. To establish whether TRIP12 and AREL1 cooperate with UBE2L3 in this process, we examined pro-IL-1β abundance following their silencing in iBMDM^YFP-UBE2L3^ cells. Indeed, silencing TRIP12 or AREL1 blocked the accelerated loss of pro-IL-1β protein seen with UBE2L3 overexpression (*Figure 4E*), which provides further evidence that these E3 ligases are involved in UBE2L3-driven enhanced turnover of pro-IL-1β. Taken together, we conclude that TRIP12 and AREL1 promote pro-IL-1β turnover and reduce the cellular pool available for caspase-1—mediated proteolytic maturation.

**Figure 4:**
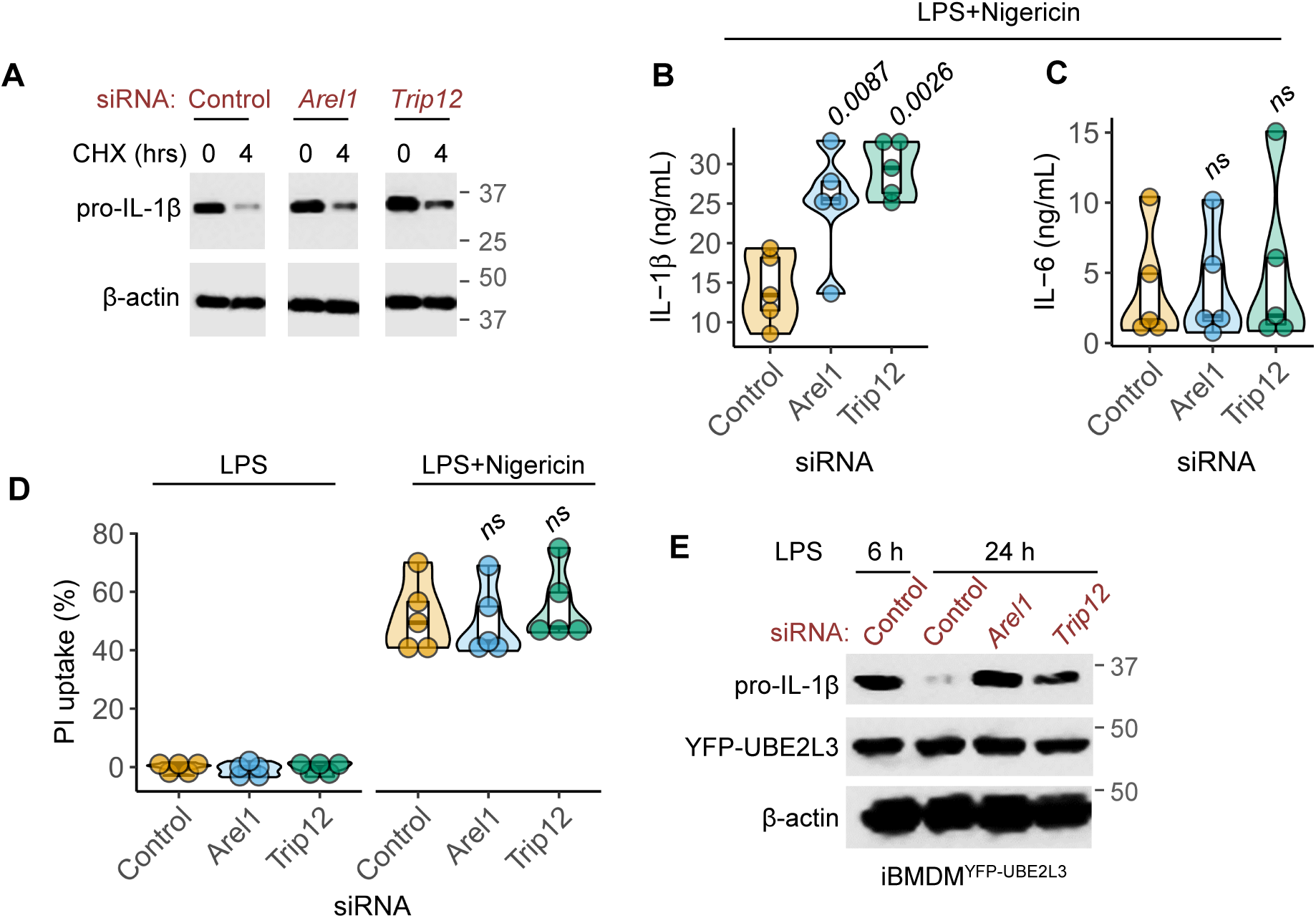
TRIP12 and AREL1 promote pro-IL-1β clearance and suppress mature IL-1β production. **(A)** Representative immunoblots from cycloheximide (CHX)-chase experiments to assess the impact of silencing AREL1 or TRIP12 on pro-IL-1β stability. iBMDMs were transfected with non-targeting Control or the indicated siRNA for 72 h, then treated with LPS (250 ng.mL^-1^) for 14 h and CHX (10 μg.mL^-1^) for the indicated time. Cell lysates were immunoblotted for pro-IL-1β and β-actin as loading control. Images from the same exposure of the same immunoblots are shown with intervening lanes removed. **(B-C)** Quantification of IL-1β **(B)** and IL-6 **(C)** by ELISA in the supernatants of iBMDMs given the indicated siRNA for 72 h, followed by inflammasome activation with LPS (250 ng.mL^-1^, 3 h) and nigericin (50 μM, 1 h). **(D)** Pyroptotic death measured as cells positive for propidium iodide (PI) dye. iBMDMs were transfected with the indicated siRNA for 72 h, followed by inflammasome activation as described in **B**. **(E)** Representative immunoblots to assess the abundance of pro-IL-1β in lysates of iBMDMs stably expressing YFP-UBE2L3. Cells were transfected with non-targeting Control or the indicated siRNA for 72 h and then treated with LPS (250 ng.mL^-1^) for the indicated times. Images in **A**, **E** represent experiments done at least three times. In **B-D** each dot represents a biologically independent experiment. Data distribution is depicted with violin, box (25th to 75th percentile, line at median), and whiskers (± 1.5 x IQR). Two-tailed *P* values for comparisons with samples given Control siRNA in **B-D** from mixed effects ANOVA. ns – not significant (*P* > 0.05).

### Pro-IL-1β is ubiquitylated by TRIP12 and AREL1

We next investigated whether pro-IL-1β interacted with AREL1 or TRIP12. Due to the lack of suitable antibodies against mouse TRIP12 and AREL1 for coimmunoprecipitation of endogenous proteins and failure of antibodies against the human proteins in these experiments, we took advantage of transient transfections in HEK293E cells to assess their interactions. These experiments showed that pro-IL-1β specifically co-immunoprecipitated with TRIP12 and AREL1 (*Figure S4A*), which indicates that these proteins can form stable complexes.

To address whether TRIP12 or AREL1 could stimulate pro-IL-1β ubiquitylation we generated HEK293E cells stably expressing pro-IL-1βHis with C-terminal hexa-histidine tag that allowed denaturing pull-downs (to remove non-covalent protein-protein interactions) with immobilised metal affinity chromatography. We co-transfected plasmids encoding HA-tagged wildtype ubiquitin along with either TRIP12 or AREL1, or mVenus as negative control, followed by denaturing pull-down and anti-HA immunoblots. This revealed marked ubiquitylation of pro-IL-1β by both E3 ligases as seen by ‘ubiquitin smears’ in western blots (*Figure 5A-B, S4B-C*).

**Figure 5.**
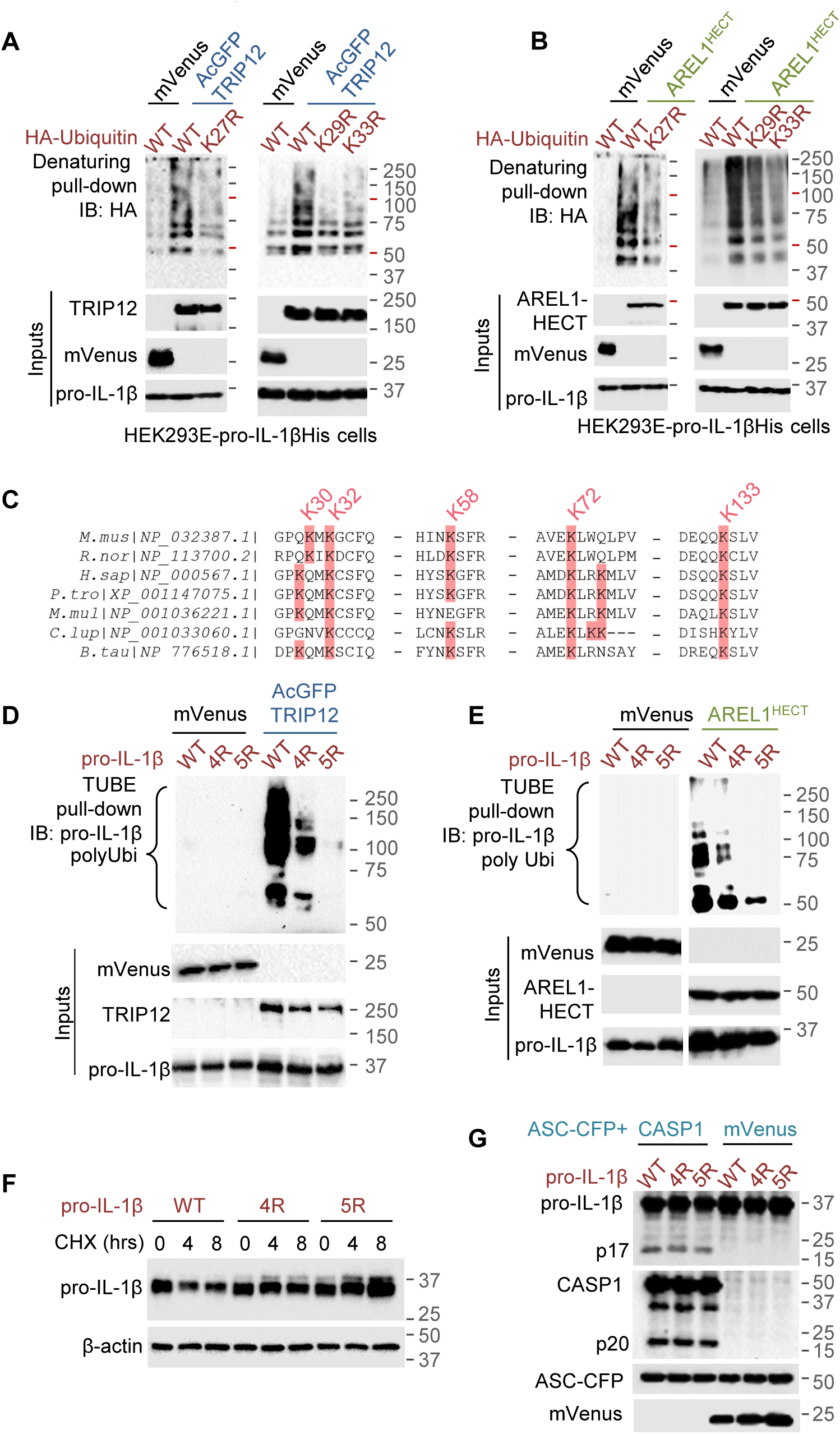
TRIP12 and AREL1 ubiquitylate pro-IL-1β in the N-terminal ‘pro’ domain. **(A-B)** Representative immunoblots from Ni-NTA pull-downs of pro-IL-1βHis under denaturing conditions (8 M urea-containing buffers) to assess the type of ubiquitin chains covalently added on pro-IL-1β by TRIP12 **(A)** or AREL1 **(B)**. HEK293E cells stably expressing pro-IL-1βHis were transfected with HA-tagged wildtype or the indicated K◊R mutants of ubiquitin along with AcGFP-TRIP12 **(A)** or AREL-HECT^Myc^ **(B)** or mVenus as negative control. Expression of proteins is shown below (Inputs). Also see *Figure S4* for additional blots from the same experiments. **(C)** Multiple sequence alignment of pro-IL-1β from the indicated organisms showing sequences flanking ‘pro’ domain lysine residues in the mouse (30, 32, 58 and 72) and human. Sequences around lysine 133 in the mature IL-1β region are also shown. *B. tau – Bos taurus, C. lup – Canis lupus familiaris, H. sap – Homo sapiens, M. mul – Macaca mulatta, M.mus – Mus musculus, P. tro – Pan troglodytes, R. nor – Rattus norvegicus.* **(D-E)** Representative images from TUBE pull-downs assays of wildtype or the indicated K◊R variants of pro-IL-1β to assess their ubiquitylation by TRIP12 **(D)** or AREL1 **(E)**. HEK293E cells were transfected with plasmids encoding the indicated pro-IL-1β variants (wildtype, 4R (K30,32,58,72R) and 5R (K30,32,58,72,133R)) and AcGFP-TRIP12 **(D)** or AREL1-HECT^Myc^ **(E)** and mVenus as negative control. Ubiquitylated proteins bound to hexa-histidine-tagged TR-TUBE were enriched with Ni-NTA beads and immunoblotted with anti-IL-1β antibody. Polyubiquitylated pro-IL-1β smears are labelled (polyUbi), and expression of proteins is shown below (Inputs). Images from the same exposure of the same immunoblots are shown in **E** with intervening lanes removed. **(F)** Representative immunoblots of cycloheximide (CHX)-chase experiments showing increased stability of the indicated variants of pro-IL-1β as compared to wildtype protein. HEK293E cells were transfected with plasmids coding for the indicated pro-IL-1β proteins, and treated with CHX (10 μg.mL^-1^) for the indicated times 18 h after transfection. **(G)** Representative immunoblots showing similar proteolytic maturation of wildtype and the indicated variants of pro-IL-1β by caspase-1. HEK293E cells were transfected with plasmids encoding wildtype or mutant pro-IL-1β without or with caspase-1 and ASC-CFP for 18 h, and cell lysates immunoblotted with the indicated antibodies. Images in **A-B**, **D-G** are representative of experiments performed at least three times.

We next asked which poly-ubiquitin chain types were added on to pro-IL-1β by TRIP12 and AREL1. To address this question, we used similar experiments as above, but with HA-tagged variants with a single K→R mutation at each lysine residue (K6R, K11R, K27R, K29R, K33R, K48R and K63R) to abrogate poly-ubiquitin chains of those types. Denaturing Ni-NTA pull-downs of pro-IL-1β^His^ immunoblotted with anti-HA antibodies revealed reduced ubiquitylation by TRIP12 with HA-Ubi^K27R^, HA-Ubi^K29R^ or HA-Ubi^K33R^ variants (*Figure 5A, S4B*), which suggested that TRIP12 adds K27-, K29- and K33-ubiquitin chains on pro-IL-1β. Similar experiments with AREL1 revealed reduced pro-IL-1β ubiquitylation in cells containing HA-Ubi^K27R^ and HA-Ubi^K33R^, indicating that AREL1 preferentially added K27- and K33-ubiquitin chains (*Figure 5B, S4C*). Taken together, we conclude that TRIP12 and AREL1 HECT-type E3 ligases target pro-IL-1β for ubiquitylation with K27-, K29- and K33-poly-ubiquitin chains.

### TRIP12 and AREL1 ubiquitylate pro-IL-1β in the ‘pro’ domain

We next asked which lysine residues in pro-IL-1β are preferentially ubiquitylated by TRIP12 and AREL1. Lys 133 in the mature cytokine region is a site for ubiquitylation [18-20], however, a K133R mutant is still ubiquitylated suggesting that other sites, including in the ‘pro’ domain, may exist [18-20]. To test whether ‘pro’ domain lysines were sites for ubiquitylation, we mutated all four lysines to arginine to generate pro-IL-1β-4R (K30R/K32R/K58R/K72R; *Figure 5C*). For these experiments we co-transfected HEK293E cells with plasmids encoding pro-IL-1β variants and either TRIP12 or AREL1 or mVenus as negative control, and enriched all ubiquitylated proteins by pull-downs with recombinant hexahistidine-tagged trypsin-resistant tandem ubiquitin interacting entities (H6-TR-TUBE, *Figure S4D*) followed by immunoblotting against pro-IL-1β (referred to as TUBE assays [21, 22]). TUBE assays revealed a drastic reduction in ubiquitylation of pro-IL-1β-4R by TRIP12 and AREL1 as compared to wildtype pro-IL-1β (*Figure 5D-E*). These results showed that the ‘pro’ domain is a major site for ubiquitylation. Consistent with ubiquitylation at K133, the additional K133R mutation in pro-IL-1β-4R (pro-IL-1β-5R) led to a further reduction in ubiquitylation (*Figure 5D-E*). We therefore conclude that the ‘pro’ domain of pro-IL-1β in addition to K133 contains major sites of ubiquitylation by TRIP12 and AREL1.

### Pro-IL-1β ubiquitylation affects its stability but not proteolysis

To assess whether the reduced ubiquitylation of pro-IL-1β mutants affected their turnover, we performed cycloheximide-chase experiments. As expected with the destabilising role of ‘pro’ domain ubiquitylation, both pro-IL-1β-4R and pro-IL-1β-5R variants were more stable than wildtype pro-IL-1β (*Figure 5F*). We next asked whether these variants could be converted to similar extent as wildtype pro-IL-1β by caspase-1. To assess their proteolytic maturation, we co-transfected them in HEK293E cells with plasmids encoding caspase-1 and ASC-CFP, which generates active caspase-1 due to spontaneous oligomerisation of CFP. Proteolysis of wildtype pro-IL-1β was similar to that of pro-IL-1β-4R and pro-IL-1β-5R (*Figure 5G*), which indicates that these lysine residues do not contribute cleavage by caspase-1. Taken together, we conclude that pro-IL-1β ubiquitylation affects its stability but not susceptibility to caspase-1—dependent proteolysis.

## DISCUSSION

We generated new *Ube2l3^ΔMac^* mice and showed that loss of UBE2L3 in macrophages/monocytes is sufficient for elevated IL-1β and inflammation following inflammasome activation. This mirrors the sufficiency of NLRP3 activation in macrophages in driving autoinflammatory disease [23], and underscores the importance of these phagocytes in IL-1β—driven pathology. Mechanistically, UBE2L3 co-opted the HECT-type E3 ligases TRIP12 and AREL1 which ubiquitylated pro-IL-1β on lysine residues mainly in the ‘pro’ domain with K27-, K29- and K33-poly-ubiquitin chains. UBE2L3, TRIP12 and AREL1 act upstream of inflammasomes and caspase-1 to impede inflammation by reducing the cellular pool of the pro-IL-1β precursor.

UBE2L3 is a highly conserved E2 enzyme (100% amino acid identity between humans and mice and 95% between humans and *Xenopus*) that participates in diverse pathways, including NF-κB signalling, p53 regulation, DNA repair, mitophagy, among others [10]. In humans, *UBE2L3* polymorphisms are linked to systemic lupus erythematosus, rheumatoid arthritis, juvenile idiopathic arthritis, inflammatory bowel disease, and ulcerative colitis [10]. Given the ubiquitous expression of UBE2L3, its cell type- and context-specific actions need to be further examined to pinpoint disease mechanisms [10]. In addition, bacterial virulence factors, such as NleL from enterohaemorrhagic *E. coli,* SopA from *Salmonella,* SidC from *Legionella pneumophila,* and OspG from *Shigella* partner with UBE2L3 during infection [24]; the antimicrobial role of UBE2L3, if any, also deserves to be explored in the future.

UBE2L3 can have exquisite specificity in its actions in cells as seen in studies on E3 ligases PARKIN and LUBAC. For instance, UBE2L3 and UBE2D1-3 serve PARKIN for the initiation of mitophagy, whereas UBE2N is required for later steps of the process [25, 26]. Similarly, UBE2L3—LUBAC partnership may be context-dependent. We found that *Ube2l3-* deletion in macrophages or its silencing with siRNA [6] does not affect NF-κB—dependent outputs (e.g., IL-6 production or *Il1b* transcription), which contrasts its role in fibroblasts [27]. As LUBAC is essential for NF-κB signalling [28], this suggests that E2 enzymes other than UBE2L3 might partner LUBAC in macrophages. Notably, the deletion of *Ube2l3* or components of LUBAC (i.e., *Rbck1* and *Rnf31*) results in embryonic lethality; whether this is due to their actions together in haematopoiesis and vascularisation during embryogenesis remains to be determined.

Pro-IL-1β ubiquitylation has been reported previously [18-20, 29], however, the E3 ligases involved were not known. We identified TRIP12 and AREL1 in an RNAi screen specifically designed to find E3 ligases that promote pro-IL-1β turnover. TRIP12 and AREL1 are ubiquitously expressed, but they have mainly been studied in relation to oncogenesis. For instance, TRIP12 plays a role in promoting DNA damage repair [30] and ubiquitin fusion degradation [31, 32], and AREL1 blocks apoptosis by targeting pro-apoptotic proteins [33], and is known to assemble K33-poly-ubiquitin chains [34-36]. Here we ascribed anti-inflammatory roles to them via their suppressive action on IL-1β production. Importantly, our identification of TRIP12 and AREL1 in pro-IL-1β turnover provides a foundation for rationally designing small molecules, e.g., PROTACs (proteasome targeting chimeras), to target IL-1β—driven inflammation. PROTACs are small molecules that leverage the ubiquitin-proteasome system for the removal of proteins of interest (POI), including neo-substrates that are not natural substrates of an E3 ligase [37]. PROTACs act as molecular glues between a POI and a convenient E3 ligase (e.g., CRBN, CRL2^VHL^), which results in POI ubiquitylation and degradation. Indeed, TRIP12 promotes the complete degradation of the oncogenic transcription factor BRD4 by its PROTAC; here the E3 ligase CRL2^VHL^ initiates ubiquitylation of BRD4, but TRIP12 adds K29- and K48-chains that enhance degradation [38]. It is tempting to speculate that TRIP12 and AREL1 could be exploited to destabilise their natural substrate pro-IL-1β using a similar approach.

K27-, K29- and K33-poly-ubiquitin chain types are associated with proteasomal degradation of several proteins. For instance, like K48-chains, K27-chains are important facilitators of substrate processing by proteasomes [39]. K27- and K29-chains destabilise several key innate immune proteins, including NLRP3 [40], MDA5 and RIGI [41, 42], ULK1 [43], STING [44], IRF3 and IRF7 [45], among others [46, 47]. Similarly, K33-chains are recognised in the turnover of RAS GTPases [48], glycine decarboxylase [49], and 3-hydroxy-3-methylglutaryl CoA reductase [49]. However, in our experiments TRIP12 and AREL1 did not modify pro-IL-1β with K48- or K11-ubiquitin chains, which have been reported on pro-IL-1β by us and others [6, 18-20]. It is therefore plausible that additional E3 ligases participate in pro-IL-1β ubiquitylation with other chain types.

Multiple ubiquitylation sites in pro-IL-1β are known. Our work complements recent findings from pro-IL-1β-K133R knock-in mice which display elevated pro-IL-1β abundance due to reduced ubiquitylation at Lys133 in the C-terminal mature cytokine region [20]. Importantly, it was noted in that study that sites other than K133 are also ubiquitylated. Our study substantially adds to these findings by identifying ubiquitylation sites in the ‘pro’ domain. The N-terminus of pro-IL-1β became of interest to us after a fortuitous finding while designing the doxycycline-inducible system for the RNAi screen. We found that N-terminal 3xFlag-tag on pro-IL-1β markedly reduced its turnover in macrophages, but C-terminal tagging had no effect (data not shown), which pointed to the importance of the native N-terminus and its intolerance to even small tags. In our study, the increased abundance of pro-IL-1β through different ways, for example, *Ube2l3-*deletion, silencing of TRIP12 or AREL1 expression, mutation of all four lysine residues in the ‘pro’ domain (pro-IL-1β-4R) or the additional mutation of K133R (pro-IL-1β-5R), all resulted in increased pro-IL-1β accumulation. The increased accumulation of pro-IL-1β led to enhanced mature IL-1β production. We therefore conclude that the entire cellular pool of pro-IL-1β is accessible to caspase-1, and the abundance of the pro-IL-1β precursor determines how much mature, inflammatory cytokine is produced. Moreover, this suggests that caspase-1 is not saturated by the levels of pro-IL-1β in wildtype macrophages and the spare proteolytic capacity is exposed by the pathological outcomes of *Ube2l3*-deletion. This highlights the importance of pro-IL-1β ubiquitylation and disposal for homeostasis.

Outstanding questions still remain, such as, which are the E3 enzymes that catalyse K63 ubiquitylation of pro-IL-1β as reported in other studies [18, 19]? Does pro-IL-1β ubiquitylation enhance its maturation [18, 19] or have no impact as reported more recently [20]? Furthermore, understanding how UBE2L3 itself is targeted by caspase-1 will shed light on the molecular circuit that governs mature IL-1β production.

Deregulated IL-1β release is linked to cancer and inflammatory conditions that affect major organs, including the vasculature, brain, liver, skin, gut, and joints [1, 2]. Currently approved therapies for neutralising IL-1 are biologics such as recombinant IL-1 receptor antagonist (anakinra), anti-IL-1 antibody (canakinumab) or a fusion protein including the ligand-binding regions of the IL-1 receptor 1 and IL1R1 accessory protein (rilonacept) [50].

However, these agents act on the cytokine already in circulation. Pro-IL-1β released from necrotic/pyroptotic cells might also be processed extracellularly [1, 2]. Targeting intracellular pro-IL-1β, for example via the proteasome, could potentially be more effective as an anti-inflammatory therapeutic intervention. Our identification of the ubiquitylation mechanisms provide a foundation for such work in the future.

## ACKNOWLEDGEMENTS

Authors would like to acknowledge funding from the MRC (MR/P022138/1, MR/T00004X/1, and MR/V030930/1 to AS). We also acknowledge the MRC-funded High-throughput Single-Cell Analysis facility (HTSCAF; MR/P028225/1) at the MRC CMBI, MRC Transgenic Facility at Imperial College London for rederiving mice, and Jessica Rowley (Manager, flow cytometry) for help with flow cytometry. CM would like to acknowledge the doctoral training programme award to MRC CMBI (MR/R502376/1). We thank Jeffery W Pollard for sharing Tg(Csf1r-cre/Esr1)1Jwp GA) mice on C57/BL6 background, and Jiri Lukas and Hiroshi Ashida for plasmids. The authors thank Sandhya Visweswariah and Gad Frankel for discussions, Teresa Thurston and David Holden for critical comments on a draft of the manuscript.

## AUTHOR CONTRIBUTIONS

Investigation: AC-P, VM, CM, TM, IG, AD, GD, MJGE, LM, MM, SW, DB, ARS; Validation: AC-P, VM, GD, MM, SW; Visualisation: VM, CM, AC-P, ARS; Writing – original draft: ARS; Writing – review and editing: ARS, VM, AC-P, DB; Supervision: ARS, VM, AC-P, DB, Software: ARS; Formal analysis: AC-P, AD, ARS; Funding acquisition: ARS; Conceptualisation: ARS.

## FIGURE LEGENDS

**Figure S1. UBE2L3 is depleted *in vivo* upon inflammasome activation.**

**(A)** IL-1β in peritoneal lavage from mice given PBS, LPS (5 μg, 3 h) or LPS+ATP (5 μg 3 h + 50 μmol, 10 min) intraperitoneally.

**(B-C)** Flow cytometry-based quantification using antibodies against F4/80 (macrophage marker) and UBE2L3 in peritoneal cells from mice given PBS, LPS or LPS+ATP as in **(A)**. Graphs show the percentage of double-positive cells **(B)** and mean fluorescence intensity in arbitrary units (AU) of UBE2L3 in F4/80+ve cells **(C)**.

**(D-)** Representative immunoblots for UBE2L3 in lysates from peritoneal macrophages from mice given PBS or LPS+ATP (5 μg for 3 h + 50 μmol 10 min) intraperitoneally as labelled. Each lane represents a mouse. Data from one of two similar experiments.

**(E)** Plot showing negative correlation between IL-1β measured in peritoneal lavage fluid and UBE2L3 staining by flow cytometry (arbitrary units, AU) in peritoneal macrophages from the same mouse. Each dot represents a mouse given PBS, LPS or LPS+ATP as in **A-C**.

Each lane in **D**, and dot in graphs represents an individual mouse. Data distribution is depicted with violin, box (25th to 75th percentile, line at median), and whiskers (± 1.5 x IQR). Two-tailed *P* value for indicated comparison from mixed model ANOVA. ns – not significant (*P* > 0.05).

**Figure S2. Generation of *Ube2l3^ΔMac^* mice.**

**(A)** Schematic depiction (left) and PCR-based screening (right) for genotyping WT (*Ube2l3^fx/fx^*) and KO (*Ube2l3^ΔMac^*) primary BMDMs given 4-hydroxytamoxifen (2 μM) for 48 h. Oligonucleotide (oligo) primer binding sites are indicated. Oligo 2 binding site is lost after Cre-mediated deletion of exon 1. Oligos 1 and 4 only generate a product under conditions of PCR if homologous recombination has occurred.

**(B)** Representative immunoblots from peritoneal macrophages isolated from mice of the indicated genotypes given tamoxifen (80 mg.mL^-1^) orally on three consecutive days.

**(C)** Relative expression of *Il1b* normalised to *Gapdh* in peritoneal macrophages isolated from WT (*Ube2l3^fx/fx^*) and KO (*Ube2l3^ΔMac^*) mice given tamoxifen. Cells were treated with LPS (250 ng.mL^-1^) for 12 h before preparing RNA for qRT-PCR.

Each lane in **B** and dot in **C** represents an individual mouse. Data distribution is depicted with violin, box (25th to 75th percentile, line at median), and whiskers (± 1.5 x IQR). Two-tailed *P* value for indicated comparison from mixed model ANOVA. ns – not significant (*P* > 0.05).

**Figure S3: Validating the siRNA screening approach to identify E3 ligases involved in pro-IL-1β clearance.**

**(A)** Representative immunoblots showing the effect of the indicated inhibitors on the abundance of LPS-induced pro-IL-1β in iBMDMs. Cells were treated with LPS (250 ng.mL^-1^) for 15 h or left untreated (UT) as controls (first two lanes, respectively). Other samples are from cells treated with LPS (250 ng.mL^-1^) for a total of 18 h where the last 3 h included the indicated inhibitors or the solvent DMSO. MG132 (proteasome inhibitor) and BAY 11-7082 (LUBAC and UBE2L3 inhibitor) treatments increased pro-IL-1β abundance as compared to DMSO or other treatments.

**(B)** (Left) Schematic showing the transcription factor rtTA3 inducing the expression of pro-IL-1βStrep in the presence of doxycycline in cells stably expressing the pLTREK-proIL-1βStrep plasmid. (Right) Representative immunoblots from lysates of pLTREK-proIL-1βStrep iBMDMs transfected with non-targeting control (Ctrl) or rtTA3 siRNA for 72 h, and then treated with doxycycline for 6 h.

**(C)** Representative immunoblots from pLTREK-proIL-1βStrep iBMDMs treated with the indicated concentrations of doxycycline for 6 h, washed and incubated for either 0 or 18 h as labelled.

**(D)** Representative immunoblots from pLTREK-proIL-1βStrep iBMDMs treated with LPS (250 ng.mL^-1^) or doxycycline (500 ng.mL-1) as indicated, followed by chase for the indicated times.

**(E)** Results from siRNA screen to identify HECT and RBR E3 ligases involved in pro-IL-1β turnover. Size of the symbols indicate mean of z* scores from three independent repeats. FDR-adjusted *P* values for comparison for each of the siRNA with the ‘15 h’ group are shown by shapes of symbols (*P* < 0.01 or *P* > 0.01); ‘high’ and ‘low’ controls are also indicated by shapes of symbols. The divergent colour scheme indicates SSMD* scores and the dotted horizontal line indicate *P* = 0.01 cut-off on a -log10() scale. Trip12 and Arel1 scored above thresholds.

**(F)** qRT-PCR showing the efficacy of silencing of *Arel1*, *Trip12* and *Herc3* transcripts. iBMDMs were transfected for 72 h with non-targeting control or siRNA against AREL1, TRIP12 or HERC3 as labelled. *Gapdh* was used as normalising control.

Images in **A-D** represent experiments performed at least three times. In **F** each dot represents a biologically independent experiment. Data distribution is depicted with violin, box (25th to 75th percentile, line at median), and whiskers (± 1.5 x IQR). P values for indicated comparisons from mixed effects ANOVA.

**Figure S4. AREL1 and TRIP12 ubiquitylate pro-IL-1β in the ‘pro’ domain.**

**(A)** Representative images of immunoprecipitation (IP) and immunoblot (IB) experiments to assess the interaction between pro-IL-1β and TRIP12 or AREL1. HEK293E cells were transfected with plasmids encoding mVenus-AREL1^Myc^ or AcGFP-TRIP12 or mVenus as negative control followed by IP with anti-GFP antibody and IB with pro-IL-1β or GFP antibodies as labelled. Expression of proteins is shown on right (Input).

**(B-C)** Representative immunoblots from Ni-NTA pull-downs of pro-IL-1βHis under denaturing conditions (8 M urea-containing buffers) to assess the type of ubiquitin chains covalently added on pro-IL-1β by TRIP12 **(B)** or AREL1 **(C)**. HEK293E cells stably expressing pro-IL-1βHis were transfected with HA-tagged wildtype or the indicated K→R mutants of ubiquitin along with AcGFP-TRIP12 **(B)** or AREL-HECT^Myc^ **(C)** or mVenus as negative control. Expression of proteins is shown below (Inputs).

**(D)** SDS-PAGE of purified recombinant hexahistidine-tagged TR-TUBE (∼2 μg) followed by Coomassie staining.

Images in **A-C** from experiments performed at least three times.

## MATERIALS AND METHODS

- **KEY RESOURCES TABLE**
- **CONTACT FOR REAGENT AND RESOURCE SHARING**
- **EXPERIMENTAL MODEL AND SUBJECT DETAILS**
  - Ethics statement
  - Generation of *Ube2l3^ΔMac^*mice
  - Mammalian Cell Culture
  - Bone-marrow derived macrophages
- **METHOD DETAILS**
  - LPS endotoxic shock models in mice
  - MSU crystal-induced peritonitis
  - Treatments of cells *in vitro*
  - Immunoblotting
  - siRNA screen and analyses
  - Quantitative RT-PCR
  - Flow cytometry
  - Enzyme-linked Immunosorbent Assays
  - Tandem Ubiquitin Binding Entities (TUBE) pull-down
  - HA-ubiquitin pull down assays
  - HEK293E cell IL-1β turnover and caspase-1 cleavage assays
  - Co-Immunoprecipitation
  - Molecular cloning
  - Recombinant protein expression and purification
  - Retroviral and Lentiviral transduction
- **QUANTIFICATION AND STATISTICAL ANALYSIS**
  - Statistical analyses of *in vivo* experiments
  - Statistical analyses of *in vitro* experiments

## CONTACT FOR REAGENT AND RESOURCE SHARING

Further information and requests for resources and reagents should be directed to Avinash Shenoy.

## EXPERIMENTAL MODEL AND SUBJECT DETAILS

### Ethics statement

All strains were backcrossed and maintained on C57BL/6 background. Mice were maintained in specific pathogen free conditions at 20-22 °C and 30-40 % humidity on 12 hours light/dark cycle. All work with mice was performed in accordance with the Animals (Scientific Procedures) Act 1986 and was approved by the local ethics review committee at Imperial College London (PIL P9A000710 holder ARS). Both male and female mice (age 2–4 months) were used for harvesting bone marrow-derived macrophages and for *in vivo* experiments.

#### Generation of Ube2l3^ΔMac^ mice

Briefly, the strategy to generate *Ube2l3-*deficient mice involved flanking exon 1 of *Ube2l3* (Gene ID 22196 on chromosome 16), which contains the translational start site, with two LoxP sites to enable its deletion by Cre-driven recombination. *Ube2l3^fx/+^* mice were generated by Taconic Biosciences on an ES cell line in the C57BL/6NTac background. These mice were crossed with mice expressing a modified oestrogen receptor fused to the Cre recombinase (icre/Esr1*) under the promoter of Csf1r (also called CD115 or MCSF1R) [13]. This results in deletion of *Ube2l3* exon 1 in macrophages and Csf1r-positive myeloid cell populations upon administration of tamoxifen to mice. Csf1R-cre/Esr1* mice backcrossed 9 generations to C57BL/6 background (B6-Tg(Csf1r-cre/Esr1)1Jwp GA) were a kind gift from Jeffery W Pollard, University of Edinburgh [13]. Mice were bred at Imperial College to generate *Ube2l3^fx/fx^cre/Esr1* mice, sperm from which were used to rederive them (MRC Transgenic Facility) into the local specific pathogen free (SPF) facility. Subsequent crosses involved homozygous *Ube2l3^fx/fx^* mice mated with similar homozygous mice carrying a single copy of Csf1r-icre/Esr1*. For convenience, mice with conditional deletion of *Ube2l3* in Csf1r-positive cells (Csf1r is highly expressed in macrophages and monocytes, with lower expression in granulocytes and lymphocytes [13, 51]) are called *Ube2l3^ΔMac^* here. All mice were genotyped at weaning (21 day old) using ear biopsies. Mice of unwanted genotypes were humanely euthanised. To induce recombination, mice were orally gavaged on 3 consecutive days with 80 mg.kg^-1^ body weight tamoxifen prepared in corn oil. Two days after the last tamoxifen treatment, *Ube2l3* deletion was confirmed by three independent methods: PCR analysis on genomic DNA from peritoneal cells using specific primers (designed by Taconic) to verify homologous recombination; flow cytometry analysis of peritoneal cells using antibodies against CD115 and UBE2L3; western blot analysis on adherent peritoneal cells using UBE2L3-specific antibody. Subsequently, all experiments were performed 48 h after the third tamoxifen dose.

#### Bone-marrow derived macrophages

Bone marrow-derived macrophage cells (BMDMs) were prepared as described before [6]. Briefly, femur, tibia, fibula were excised from mice, cleaned in 70 % ethanol and DMEM containing gentamicin (50 μg.mL^-1^), and cells in the marrow flushed out with 5 mL DMEM. Single-cell suspensions were generated by vigorous pipetting and used for differentiation into macrophages or stored in 90% heat-inactivated foetal bovine serum (HI-FBS) containing 10 % DMSO for future use. Differentiation was carried out in non-cell culture-treated (bacterial) 10 cm petri plates in medium containing 20 % conditioned-medium from L929 fibroblast culture. Macrophages were used for experiments after day 6 of differentiation. BMDMs were immortalized using J2CRE virus as described before to generate immortalized BMDMs (iBMDMs) and reduce the use of mice [6, 52]. iBMDMs were maintained in Dulbecco’s modified Eagle’s medium (DMEM) supplemented with 10% HI-FBS, 100 U.mL^-1^ penicillin and streptomycin, and 20% L929 conditioned-medium.

#### Mammalian Cell Culture

Primary BMDMs and iBMDMs were grown in complete DMEM (high-glucose DMEM, 10% HI-FBS, 50 U.ml^-1^ penicillin and 50 µg.ml^-1^ streptomycin) supplemented with 20% L929 conditioned-medium. Additionally, puromycin (6 µg.ml^-1^), doxycycline (500 ng.mL^-1^) were added when needed. Human embryonic kidney 293E (HEK293E) and L929 cells were grown in complete DMEM medium. Cells were routinely tested, and found negative, for mycoplasma contamination.

## METHOD DETAILS

### LPS endotoxic shock models in mice

Mice were treated with 2.5 mL of 3% sterile thioglycolate intraperitoneally to induce peritoneal macrophages. Two days after thioglycolate injections, *Ube2l3* deletion was induced in mice by oral administration of tamoxifen as described above. For the high-dose model, mice were given 30 mg.kg^-1^ body weight of LPS prepared in sterile PBS for 3 h intraperitoneally. Disease severity was measured as the sum of four parameters: coat fur, mobility, bulging of eyes and posture; scores ranged from 0 to 4, wherein 0 scores for a healthy mouse. Scoring was done by a trained professional blinded to treatment and genotype of mice. Mice were euthanised and blood was collected for serum cytokine analyses.

The low-dose LPS model involved thioglycolate and tamoxifen treatments as above. LPS (5 μg in sterile PBS) was given intraperitoneally for 3 h followed by an intraperitoneal injection of 50 μmol of ATP (in sterile PBS). Ten minutes after ATP treatment, mice were euthanised, blood and peritoneal lavage were collected for cytokine analyses. Peritoneal lavage was collected by injecting 3 mL sterile ice-cold Dulbecco’s modified PBS containing 0.1 % BSA.

For *Ube2l3* depletion post inflammasome activation, *in vivo* peritonitis was induced in thioglycolate treated *Ube2l3^fx/fx^* mice as described above, and peritoneal lavage was collected, and 1 × 10^6^ cells were stained for flow cytometry analysis as described below and supernatant was assayed for cytokines using manufacturer’s protocol.

### MSU crystal-induced peritonitis

MSU-induced peritonitis was performed as described previously [4, 53]. In brief, following tamoxifen regimen as above, adult mice were injected intra-peritoneally with 1 mg MSU crystals resuspended in 0.2 mL PBS. After 6 h, mice were euthanized and injected with 1 mL PBS into the peritoneum. Peritoneal lavage was collected, and 1 × 10^6^ cells were stained for flow cytometry analysis and supernatant was assayed for IL-1β using manufacturer’s protocol.

### Treatments of cells in vitro

To induce *Ube2l3* knock-out in primary or immortalized *Ube2l3^fx/fx^* and *Ube2l3^fx/fx^ ^iCre+^* BMDMs in 10 cm dishes were treated with 2 μM of 4-hydroxytamoxifen (hTam) for 48 h, followed by trypsinisation to harvest cells which were then plated for experiments as required.

For cycloheximide (CHX)-chase experiments, hTam-treated cells or siRNA transfected iBMDMs were plated in 48 well plate at a seeding density of 1.5 x 10^5^ cells per well. Cells were treated with 250 ng.mL^-1^ of LPS for 14 h followed by treatment with 10 μg.mL^-1^ of CHX for times as indicated. Inhibition of proteasomes was achieved by treating cells with 10 μM of MG-132.

To determine the effect of various chemical inhibitors of protein degradation pathways on pro-IL-1β abundance, cells were primed with LPS (250 ng.mL^-1^) or doxycycline (500 ng.mL^-1^) and followed by treatment with MG-132 (10-25 µM), Bafilomycin A (BafA; 20 nM), BAY 11-7082 (50 μg.mL^-1^), MLN4924 (3 µM) and Heclin (100 µM) at times as indicated in figure legends.

To determine the levels of mature IL-1β in iBMDMs, cells were treated with 250 ng.mL^-1^ LPS for 3 h, followed by 50 μM of nigericin for 1 h. For western blots, samples were treated and processed as described before [6]. Secreted IL-1β was measured using ELISA and western blot analysis of supernatant from cells. Cell cytotoxicity was measured using propidium iodide (PI) dye uptake assay [54]. Cells lysed with Triton-x100 was used to obtain the percentage of the PI uptake in treated cells and untreated cells were used as baseline controls.

### Immunoblotting

Samples for immunoblotting were prepared in RIPA buffer (60 mM Tris PH 8.0, 150 mM NaCl, 1 % NP-40, 0.5 % sodium deoxycholate, 1 mM EDTA) supplemented with 1x protease inhibitor and 1 mM PMSF, and then mixed with Laemmli loading buffer (50 mM Tris pH 6.8, 2% (w/v) SDS, 10% (v/v) glycerol, 0.01% (w/v) bromophenol blue, 0.05% (w/v) 2-mercaptoethanol) [6, 54, 55]. Extracts were separated by SDS-PAGE using Tris-Glycine buffer systems and transferred to PVDF membranes. Blots were incubated with the indicated primary and secondary antibodies; polyclonal anti-UBE2L3 (GTX104717; GeneTex), goat anti-mouse IL-1β (AF401; R&D systems), anti-FLAG M2 (Sigma-Aldrich), Anti-GFP (sc-9996, Santa Cruz), anti-β-actin-HRP (A3854; Sigma-Aldrich), anti-Myc (sc-40, Santa Cruz). Immunoblots were developed with Clarity Western ECL for cell lysates and ECL Prime for supernatant samples. In all images, molecular weights markers are marked in kDa units based on migration of Precision Plus Protein Dual Color Standards.

### siRNA screen and analyses

To identify the E3-ligase involved in the ubiquitylation of pro-IL-1β, we used iBMDMs expressing pro-IL-1β under doxycycline-inducible promoter coded in the pLTREK-2P-mIL-1βStrep vector (derived from a vector described previously [55]). A cherry-picked library of SMART Pool siRNA for mouse HECT and RBR ubiquitin E3 ligases and non-targeting controls was obtained from Dharmacon™ where genes were randomly assigned to wells in a 96-well plate, including empty wells, non-targeting controls, *rtta3* siRNA. The screen was performed three independent times with technical duplicates within each repeat. Experimenters and analysts were blinded to gene names until data analyses.

Briefly, iBMDMs containing pLTREK-2P-mIL-1βStrep were seeded in a 96-well plate at a density of 3 x 10^4^ cells/well 24 h before siRNA transfection. Transfections were performed with TransIT-X2® Dynamic Delivery System (25 nM final concentration of siRNA in wells) for 72 h according to manufacturer’s procedures. Cell culture medium was changed 48 h after transfection, and treatments were performed the following day. Cells were treated with doxycycline (500 ng.mL^-1^) for 15 h, washed twice and chased for 3 h for turnover (total 18 h treatment). “High” controls (control siRNA + 3h MG-132 treatment (to block proteasomal turnover) or 15 h doxycycline treatment (no ‘chase’)) and “low”controls (no doxycycline treatment, *rtta3* gene silencing, 15 h doxycycline treatment + 3 h ‘chase’) were done in wells randomly placed in the 96-well plate. After treatment, cells were lysed with PBS, 0.1%Tween-20, protease inhibitors and 1% BSA followed by 2 cycles of freeze thaw. Pro-IL-1β levels were measured by ELISA (88-8014-22; Thermo Fisher), and cell death was determined by LDH in the supernatant using Promega LDH release assay kit. Each siRNA screen attempt included technical duplicate transfections, and means from three independent repeats were analysed in Microsoft Excel and R, and plotted in R. Robust z score (z*), robust strictly standardized mean difference (SSMD*) and *P* value cut-offs were calculated following published methods [17]. ‘Robust’ z scores use medians and median of the absolute deviation (MAD) and are less affected by outliers than mean and standard deviation. Briefly, log-transformed value of pro-IL-1β in the well that received non-targeting control siRNA (‘low control’) was subtracted from log-transformed values for all other wells followed by calculation of z* scores for all siRNA target wells within each biological repeat. SSMD* scores were calculated from z* from three independently repeated screens. Mean z* from three experiments and SSMD* are plotted in Figures 3D and S3E. *P* values were calculated following linear mixed-effects analyses (random intercepts allowed for each experiment) and false discovery rate (FDR)-adjustment for comparisons for each siRNA against ‘low control’, i.e., non-targeting control siRNA-treated wells incubated up to 18 h for pro-IL-1β turnover. Genes whose silencing led to increased pro-IL-1β abundance were expected to have a low *P* value and relatively high z* and SSMD* scores.

### Quantitative RT-PCR

RNA extraction was performed using the RNeasy mini kit according to the manufacturer’s protocol. Reverse transcription used 2 µg of purified RNA and High-capacity cDNA Reverse Transcription kit. Quantitative PCR was performed using SsoAdvanced Universal SYBR green supermix on a StepOnePlus Real-Time PCR system. Reactions were performed in duplicate, including negative control lacking cDNA or primer. Data are expressed as 2^-ΔCT^ values normalised to *Gapdh*.

### Flow cytometry

Cells (1 × 10^6^) collected from the peritoneal lavage were washed in flow cytometry staining buffer and incubated with 1:100 dilution of anti-mouse CD16/32 (Fc block) antibody for 10 min at room temperature. Cell surface marker staining was performed using 1:100 dilution of respective antibodies (e.g., anti-mouse F4/80-APC, anti-mouse Ly6G-APC, or anti-mouse CD115 (Csf1r)-APC) in dark for 30 min at 4°C in staining buffer. Cells were washed thrice with staining buffer followed by fixation with IC fixation buffer for 15 min. For intracellular antigen staining, cells were washed thrice with permeabilization buffer and incubated with permeabilization buffer for 10 min at 4°C. UBE2L3 was stained using anti-mouse Ube2l3-FITC antibody at 1:100 dilution in dark for 30 min at 4°C in permeabilization buffer, followed by three washes in staining buffer. Total neutrophils were estimated by staining cells with anti-mouse Cd11b-FITC and anti-mouse Ly-6G-APC. Data was collected on a BD FACSCalibur^TM^ and analysed with Cyflogic software.

### Enzyme-linked Immunosorbent Assays

Cytokines were detected by ELISA from serum and peritoneal lavage samples of mice, and the supernatant of treated cells or cell lysates. The following ELISA kits were used to determine cytokine concentration according to the manufacturer’s instructions: mouse pro-IL-1β (88-8014-22; Thermo Fisher), mouse IL-1β (88-7013-88; Thermo Fisher), mouse TNFα (88-7324; Thermo Fisher), mouse IL-6 (88-7064; Thermo Fisher).

### Recombinant TR-TUBE expression and purification

BL21-CodonPlus (DE3)-RIPL cells were transformed with pPRO_Ex-HIST-TEV_6xTR-TUBE, and protein expression was induced with 100 μM IPTG at 18 °C for 18 h. Cells were collected, washed in ice cold PBS and lysed by sonication in lysis buffer (50 mM Tris-HCl pH 7.0, 300 mM NaCl, 15 mM imidazole, 5 mM 2-ME and 1 mM PMSF). Initial Ni^2+^-NTA purification was carried out using a HisTrap HP column (5 mL column, GE Healthcare) on an ÄKTA Start, with all buffers being ice-cold and filtered through a 0.2 μm membrane. Unbound material was washed with 10 column volumes (CV) of wash buffer (50 mM Tris-HCl pH 7.0, 100 mM NaCl, 15 mM imidazole and 5 mM 2-ME) followed by a 10 CV gradient elution over 50 fractions in elution buffer (50 mM Tris-HCl pH 7.0, 100 mM NaCl, 200 mM imidazole, 5 mM 2-ME). Fractions containing 6H-TR-TUBE (theoretical pI 3.69) were diluted in zero salt buffer (50 mM Tris-HCl pH 7.0, 5 mM 2-ME) to a final NaCl concentration of 10 mM. The sample was then loaded onto a HiTrap Q HP anion exchange column (5 mL, GE Healthcare), followed by washing unbound material using 10 CV of AEX wash buffer (50 mM Tris-HCl pH 7.0, 10 mM NaCl, 5 mM 2-ME) and elution over a 10 CV gradient of AEX elution buffer (50 mM Tris-HCl pH 7.0, 1 M NaCl, 5 mM 2-ME). Fractions containing 6H-TR-TUBE were pooled before concentration and desalting into storage buffer (50 mM Tris-HCl pH 7.0, 100 mM NaCl, 5% glycerol, 5 mM 2-ME) through two centrifugation steps using a Vivaspin 20 MWCO 5 kDa Centrifugal Filter Column. The protein concentration was estimated using the Bradford assay, and recombinant 6H-TR-TUBE was snap frozen in storage buffer and stored at -80 °C until use, with ∼20 μg protein used per pull-down.

### Tandem Ubiquitin Binding Entities (TUBE) pull-down

HEK293E cells (4 x 10^5^) were plated in 6-well plates and transfected with either wildtype or mutant mouse IL-1β along with either plasmid encoding Arel1-Hect^MYC^ (HECT domain of Arel1), AcGFP-Trip12 or mVenus as negative control. One day post-transfection, cells were treated with 10 μM MG132 for 3 h, washed with ice-cold PBS and harvested in 500 μL of lysis buffer (25 mM Tris pH 7.0, 150 mM NaCl, 1% NP-40, 20 mM imidazole, 10 mM freshly prepared N-ethyl maleimide (NEM)) containing 1 mM PMSF, and protease and phosphatase inhibitor cocktails. Harvested cells were incubated for 20 minutes on an end-over rocker at 4 °C, followed by 3 cycles of 10 seconds each of sonication at 30% amplitude (Sonics, VibraCell). Lysates were centrifuged (14,000 x*g*, 30 min, 4 °C) and supernatants incubated with TR-TUBE overnight at 4 °C on an end-over rocker. The TR-TUBE protein along with ubiquitylated proteins was pull-down using Ni-NTA magnetic beads for 1 h at 4 °C, followed by four washes in lysis buffer, and elution of bound proteins in reducing and denaturing Laemmli loading buffer for western blot analysis.

### HA-ubiquitin pull down assays

HEK293E cells with stable expression of hexa-histidine-tagged mouse pro-IL-1β were generated using retroviral transduction of HEK293E cells with pMX-CMV-mIL-1β-His plasmid, followed by selection of transduced cells with 2 μg/mL of puromycin. His-tagged mouse IL-1β expression in selected cells was confirmed using western blot analysis. HEK293E-mIL-1β-His cells (4 x 10^5^) were plated in 6-well plates and transfected with either plasmid encoding HA-tagged wildtype ubiquitin or HA-tagged mutant ubiquitin with either plasmid encoding Arel1-HECT^MYC^, AcGFP-Trip12 or mVenus as negative control. At 24 h post-transfection, cells were treated with 10 μM of MG132 for 3 h, washed with ice-cold PBS and harvested in 500 μL of denaturing lysis buffer (25 mM Tris pH 7.0, 300 mM NaCl, 1 % NP-40, 15 mM imidazole, 8 M Urea, 10 mM NEM) containing 1 mM PMSF, and protease and phosphatase inhibitor cocktails. Harvested cells were incubated 20 minutes on an end-over rocker at 4 °C, followed by 3 cycles of 10 seconds each of sonication at 30% amplitude (Sonics, VibraCell). Lysates were centrifuged (14,000 x*g*, 30 min, 4 °C) and supernatants incubated with magnetic Ni-NTA beads overnight at 4°C on an end-over rocker. Following four washes in denaturing lysis buffer, bound proteins were eluted using reducing and denaturing Laemmli loading buffer for western blot analysis using anti-HA antibody.

### Pro-IL-1β turnover and caspase-1 cleavage assays in HEK293E cells

For wildtype, 4R and 5R pro-IL-1β^strep^ turnover assays, HEK293E cells (4 x 10^5^) were plated in 6-well plates and transfected with plasmid encoding the corresponding variant. One day post-transfection, cells were harvested with trypsin and plated in five wells of a 24-well plate and left overnight. Cells were treated with cycloheximide (10 μg.mL^-1^) for times as indicated in figures and total cell lysates were prepared in RIPA buffer as above for immunoblot analysis. For caspase-1 cleavage, HEK293E cells (1 x 10^5^) were plated in 24-well plates and transfected with plasmid encoding either wildtype or mutant mouse pro-IL-1β, mASC-CFP and either mouse caspase-1 or mVenus as negative control. Cell lysates were prepared 18 h post-transfection for immunoblot analysis.

### Co-Immunoprecipitation

HEK293E cells (4 x 10^5^) were plated in 6 well plates and transfected with wildtype mouse IL-1β along with either plasmid encoding mVenus-tagged Arel1, AcGFP-tagged Trip12 or plasmid encoding mVenus as negative control. One day post-transfection, cells were treated with 10 μM MG132 for 3 h, washed with ice-cold PBS and harvested in RIPA buffer containing 1 mM PMSF, and protease and phosphatase inhibitor cocktails. Harvested cells were lysed on an end-over rocker at 4 °C for 20 minutes, followed by 3 cycles of 10 seconds each of sonication at 30% amplitude (Sonics, VibraCell). Lysates were centrifuged (14,000 x*g*, 30 min, 4 °C), and supernatants used for immunoprecipitation with 1 μg of anti-GFP antibody. Samples were incubated on an end-over rocker for 18 h at 4°C, followed by 2 h incubation with 40 μl of magnetic protein G-Sepharose slurry to capture antigen-antibody complexes. Beads were washed thrice with RIPA lysis buffer; each wash was carried out for 10 min on an end-over rocker at 4 °C. Protein complexes bound to the beads were eluted with 50 μl of 2× Laemmli loading buffer and analysed by western blotting.

### Molecular cloning

Molecular cloning was performed using one-step sequence and ligation independent cloning (SLIC) [56], and all constructs were confirmed by DNA sequencing (GATC Biotech or Genewiz). Doxycycline-inducible expression used an in-house one-step lentiviral plasmid like pLTREK3-TEV-T2-GFP used previously [55]. Briefly, mouse IL-1β cDNA was cloned downstream of the Tet-inducible promoter from pRetroXTight (Takara), with a Kozac sequence and C-terminal 2x StrepTag II (WSHPQFEKGGGSGGGSGGGSWSHPQFEK) tags. The TetOn transcription factor rtTA3 gene, a self-cleaving P2A peptide (GSGATNFSLLKQAGDVEENPGP) and puromycin-resistance gene were cloned downstream of the PGK promoter to obtain pLTREK-2P-mIL-1βStrep. pMxCMV-YFP and pMxCMV-YFP-UBE2L3 plasmids were described before [6]. Similarly, pro-IL-1β constructs with 2x C-terminal StrepTag II tags were cloned into pMX-CMV plasmid for constitutive expression.

Mouse AREL1 E3 ligase cDNA obtained from Origene (MR210828) and the different constructs (HECT-domain: aa 436-823) were cloned with two C-terminal 2x-Myc tags or N-terminal mVenus and C-terminal 2x-Myc tags into pMX-CMV-mVenus-2Myc. A T424I polymorphism in the commercial cDNA was corrected to generate I424T AREL1 for subsequent cloning. Human TRIP12 plasmid (pAcGFP-TRIP12) was a kind gift from Jiri Lucas (University of Copenhagen) [57].

Site-directed mutagenesis used single oligonucleotide mutagenesis based linear PCR as described previously [58]. In some cases, two mutagenic primers with ∼18-bp identical ends (for SLIC cloning) were used to amplify the entire plasmid by PCR with the hot-start KOD polymerase followed by SLIC [56]. This used KOD polymerase buffer containing 2.25 mM MgSO_4_ and 2.5 % DMS). Cycling conditions used two-stage programmes with first 10 cycles used annealing temperatures starting at 60 °C with 0.5 °C increase per cycle, followed by 10 cycles of 2-step PCR consisting of denaturation (94 °C) and amplification (72 °C) without an annealing step. This method was used to generate the following mutations in pMXCMV-pro-IL-1β^strep^: K30R, K32R, K58R, K72R, and K133R, and their combinations to generate pMXCMV-pro-IL-1β^strep^-4R (K30R, K32R, K58R, K72R) and pMXCMV-pro-IL-1β^strep^-5R (K30R, K32R, K58R, K72R, K133R).

TR-TUBE (4 tandem repeats of the trypsin-resistant UBQLN1 UBA domain tagged with N-terminal hexahisitine tag) from pRSET-6xTR-TUBE (Addgene, #110313, kind gift from Yasushi Saeki, [22]) was digested by BamHI and SalI enzymes and cloned into pPRO-Ex-HT-A vector to obtain pPRO_Ex-HIST-TEV_6xTR-TUBE.

### Retroviral and Lentiviral transduction

For retro-(e.g., pMX-CMV plasmids) and lenti- (e.g., pLTREK plasmids) viral transduction, plasmids were transfected with Lipofectamine 2000 into HEK293E cells plated in 12-well plates for packaging pseudotyped virus-like particles. A total of 1.5 ug DNA per well consisting of target plasmid: pCMV-MMLV GagPol (for retroviral vectors): pCMV-VSV-G at a ratio of 5:4:1 or target plasmid: pHIV GagPol (for lentiviral vectors): pCMV-VSV-G at a ratio of 4:3:2, was used. Culture supernatants were collected after 48 h after addition of sterile cell culture-grade Hepes buffer (pH 7.4, final concentration 10 mM) and filter-sterilised (0.45 µm non-PVDF 13 mm sterile filters) for transduction. Target cells plated in one well of a 12-well plate in 1 mL medium received ∼0.3-0.4 mL of packaged virus for 48 h, followed by removal of medium and addition of fresh medium containing the appropriate selection antibiotic.

## QUANTIFICATION AND STATISTICAL ANALYSIS

### Statistical analysis and data plotting of in vivo experiments

Experiments were planned as randomized blocks with 2-3 animals per genotype per experiment, and experiments were repeated independently 3-5 times. Mice were genotyped for the presence of *Ube2l3^fx/fx^* without or with Csf1R-Cre-ERT2 gene (to be able to selectively delete *Ube2l3* in macrophages) and randomly assigned to tamoxifen or oil groups, and were genotyped again after sacrifice to independently verify the presence or absence of the *Cre* gene. For high-dose LPS model, disease activity scores were determined by an experimenter blinded to genotypes and treatments. Investigators were not blinded to treatments in other experiments. Age- and sex-matched male and female mice of age between 8-12 weeks were used across experiments. Data from all mice in all experiments were pooled and analysed using mixed effects models with ‘Experiment’ as blocking factor (variable intercepts) using lme4 [59], lmerTest [60], and emmeans [61] packages as implemented in the grafify [62] package in R. When model diagnostics (e.g., ggResidpanel [63] and performance [64] packages in R) revealed major deviation of residuals from the normal distribution, data transformations (e.g., logarithms, logit) were used. False discovery rate (FDR; Q = 5 %) was used to correct for multiple comparisons as implemented in emmeans in R. FDR-adjusted *P* values >0.05 were considered non-significant (ns). Graphs were generated in R using grafify [62] and ggplot2 [65] packages, using dotplot, boxplot and violin geometries. Dotplots (default binwidth 1/30 of Y axis) are shown with data distribution depicted by a box (showing 1^st^ and 3^rd^ quartiles), with horizontal line at median and whiskers (± 1.5 interquartile range (IQR)), and violins (trimmed to minimum and maximum data values).

### Statistical analysis and data plotting of in vitro experiments

When performing the siRNA screen, experiments were blinded to gene names in wells until the analyses of independent repeats. *In vitro* experiments, such as ELISA, LDH-release, PI-uptake, real-time qRT-PCR, and others were set up as two to three technical replicate wells, values from which were averaged to obtain a mean for that experiment. Experiments were independently repeated with fresh source/passage of cells on different days (indicated by *n* in Figure Legends) and means from different experiments were analysed statistically using mixed effects models with ‘Experiment’ as random factor (variable intercept models). When model diagnostics (e.g., ggResidpanel [63] and performance [64] packages in R) revealed major deviation of residuals from the normal distribution, data transformations (e.g., logarithms, logit) were used. False discovery rate (FDR; Q = 5 %) was used to correct for multiple comparisons as implemented in emmeans in R. Graphs were generated in R using grafify [62] and ggplot2 [65] packages, using dotplot, boxplot and violin geometries. Dotplots (default binwidth 1/30 of Y axis) are shown with data distribution depicted by a box (showing 1^st^ and 3^rd^ quartiles), with horizontal line at median and whiskers (± 1.5 interquartile range (IQR)), and violins (trimmed to minimum and maximum data values).

## KEY RESOURCES TABLE

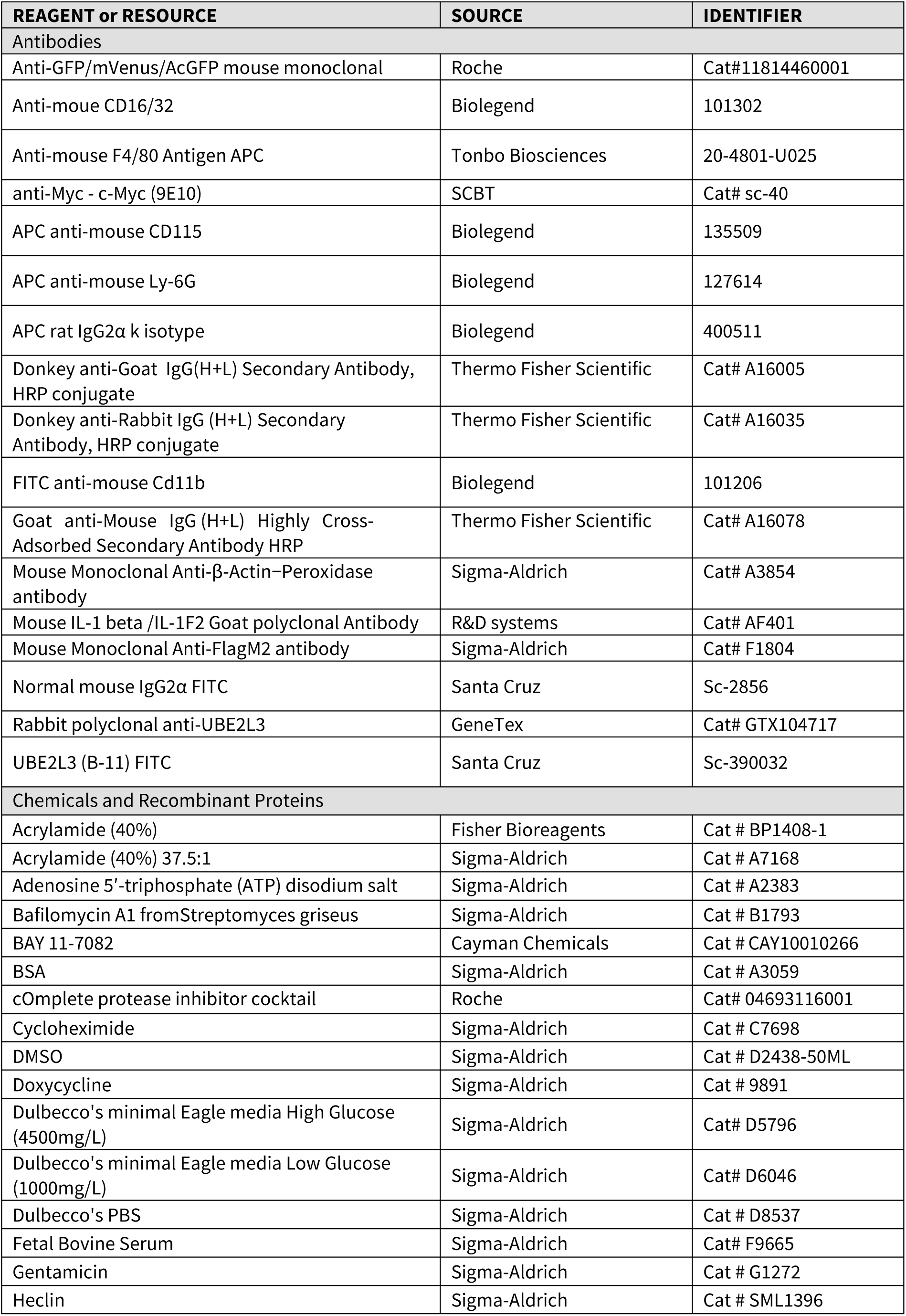
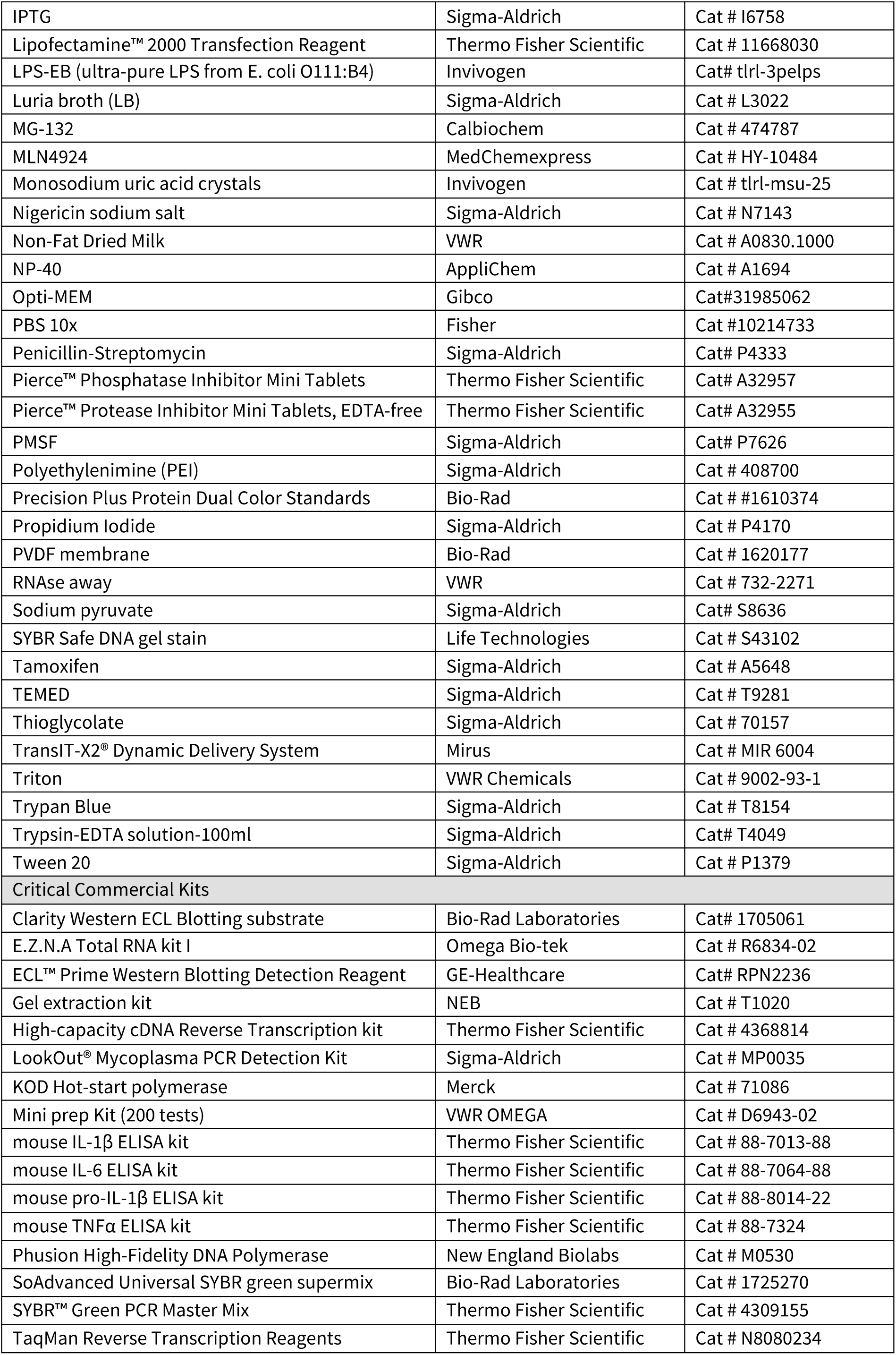
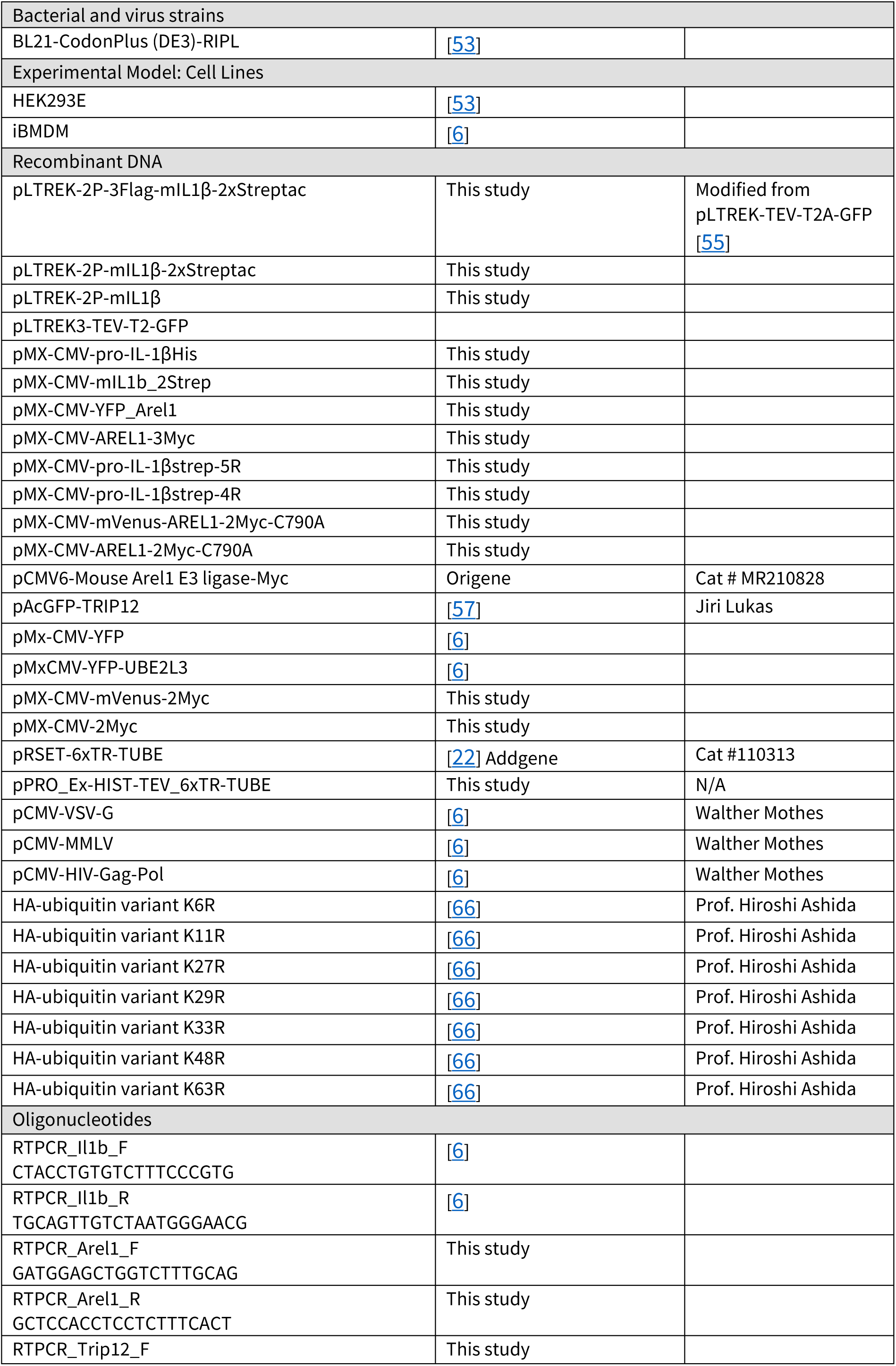
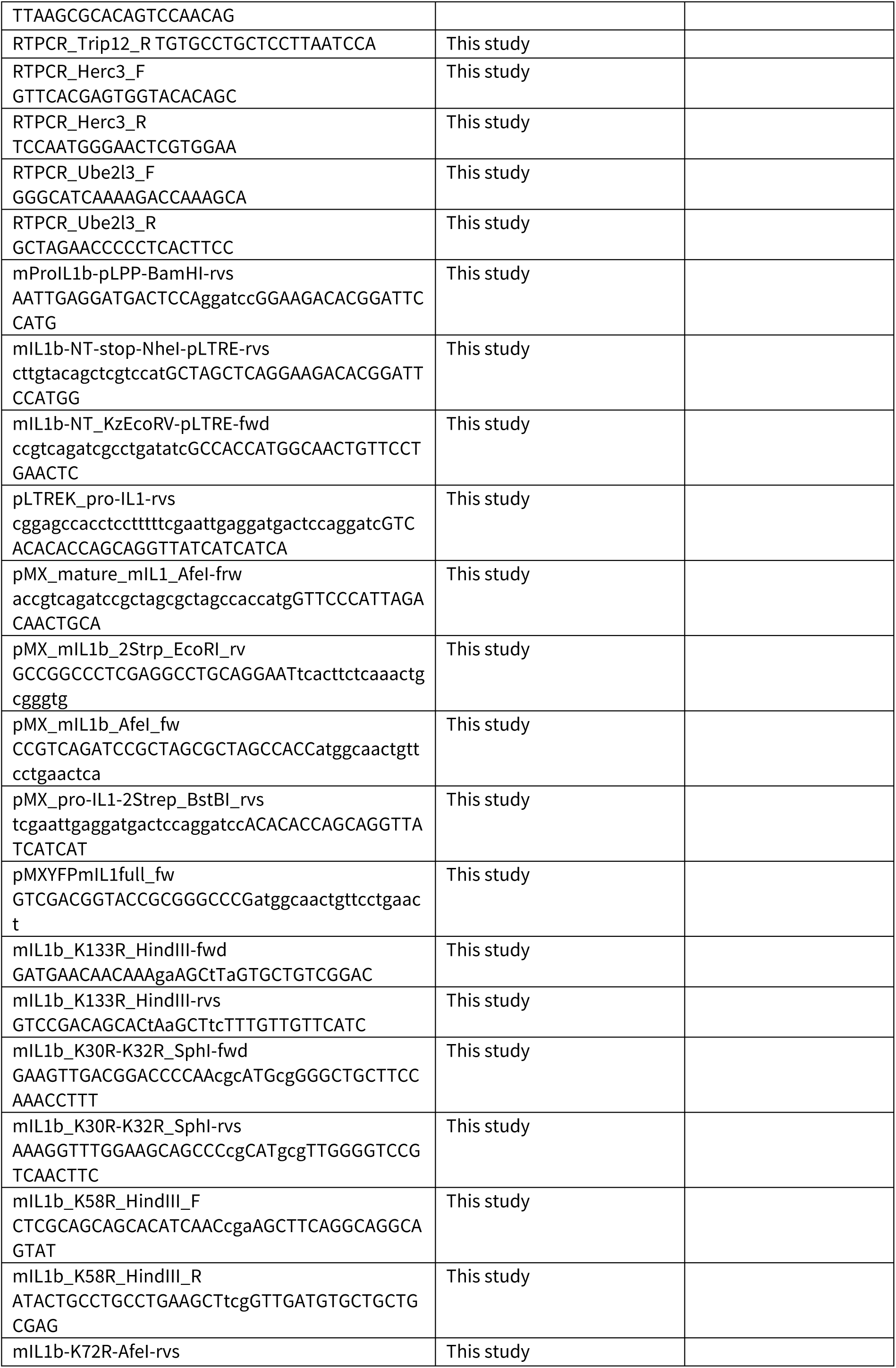
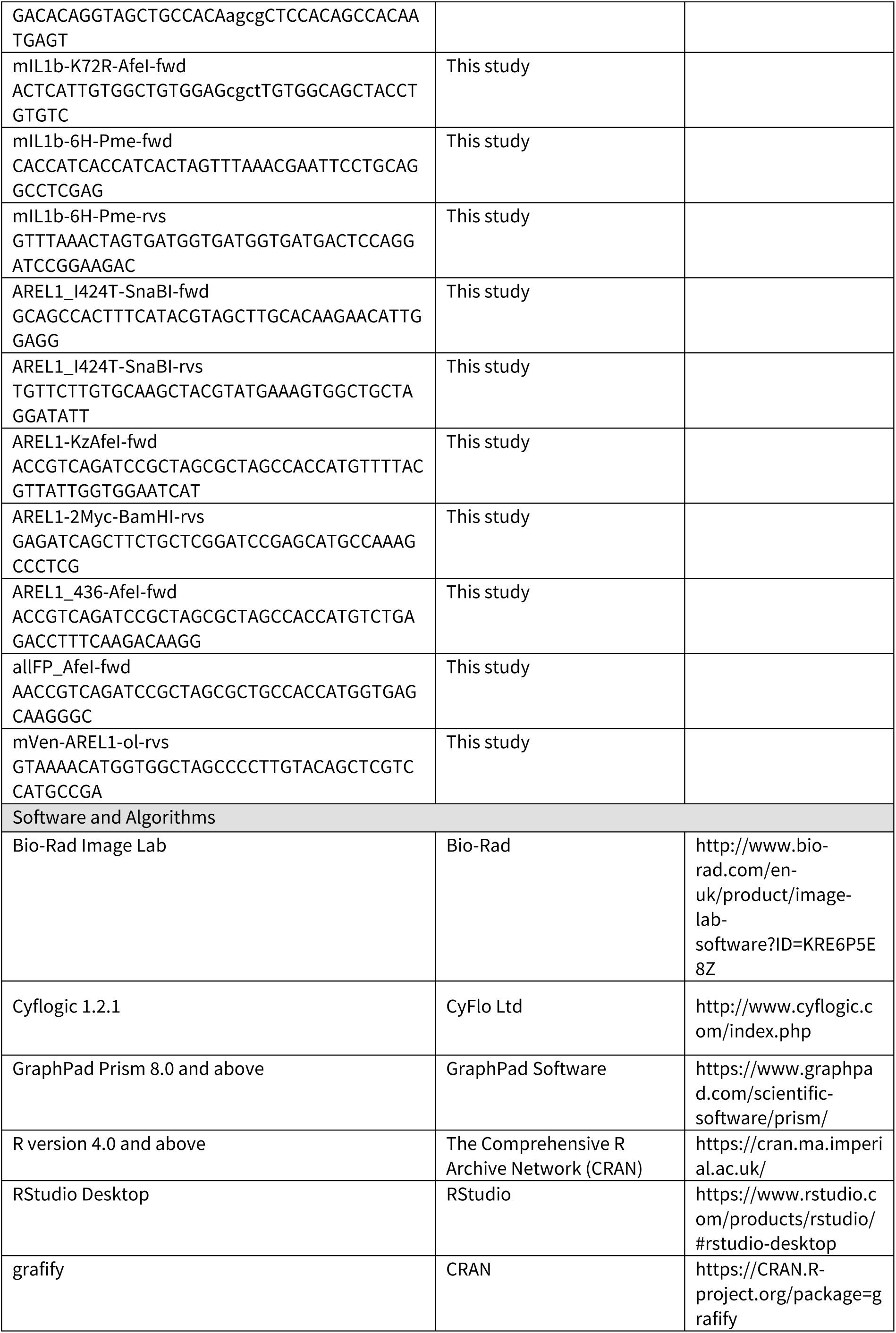
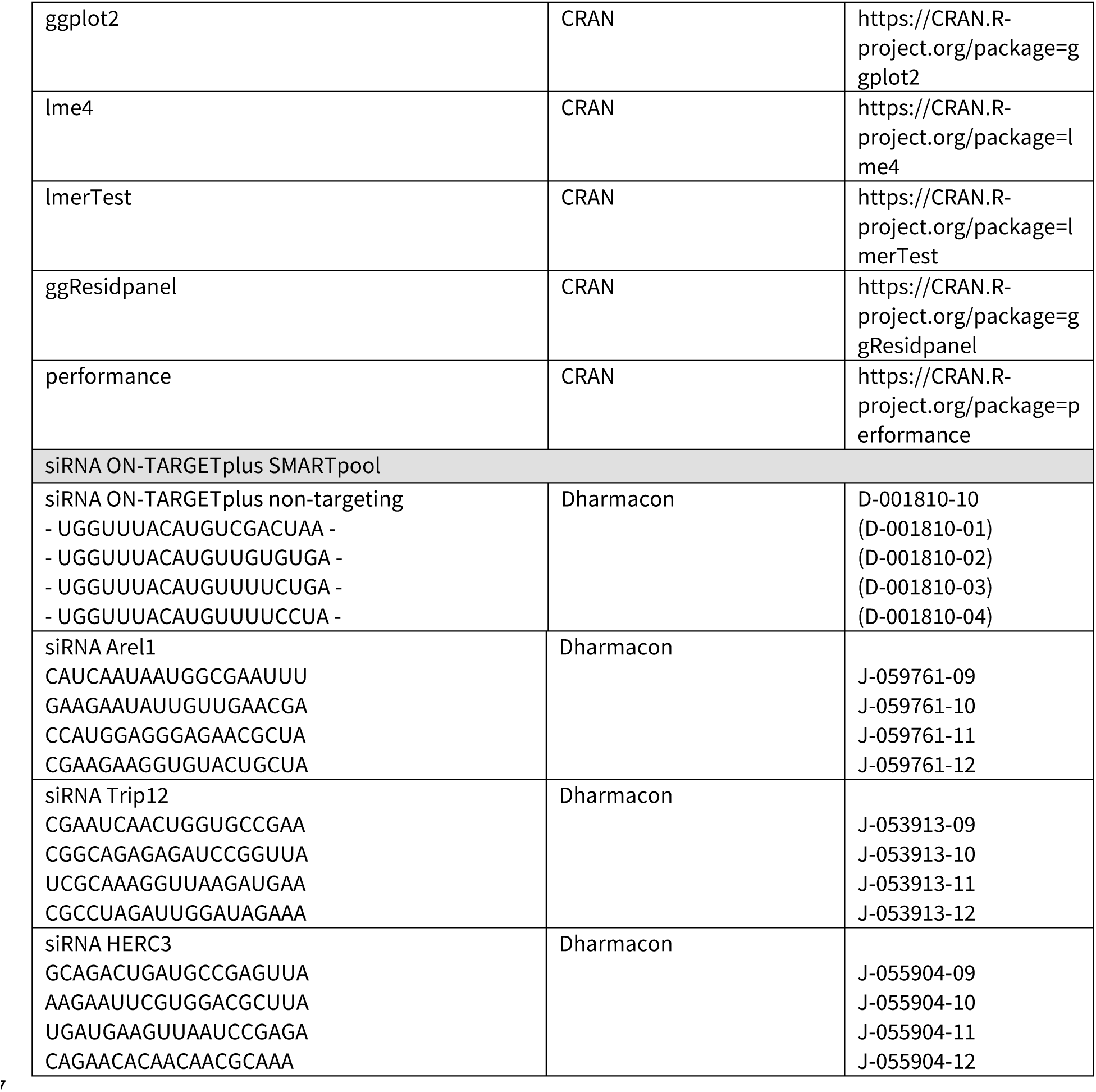

