## Supplemental Figures S1-S4 for "IL-1β turnover by TRIP12 and AREL1 ubiquitin ligases and UBE2L3 limits inflammation"

### IL-1β turnover by TRIP12 and AREL1 ubiquitin ligases and UBE2L3 limits inflammation

Vishwas Mishra[1, 2], Anna Crespo-Puig[1, 2], Callum McCarthy[1], Tereza Masonou[1], Izabela Glegola-Madejska[2], Alice Dejoux[1], Gabriella Dow[1], Matthew J. G. Eldridge[1], Luciano H. Marinelli[1], Meihan Meng[1], Shijie Wang[1], Daniel J. Bennison[1], Avinash R. Shenoy[1,*]

[1] Medical Research Council Centre for Molecular Bacteriology & Infection, Imperial College London, London, UK

[2] equal contribution

* Correspondence & Lead Contact:

Address: Room 4.40A, Flowers Bldg, Armstrong Road, Imperial College London, London SW7 2AZ, UK

**Figure S1:** *Mishra, Crespo-Puig et al*

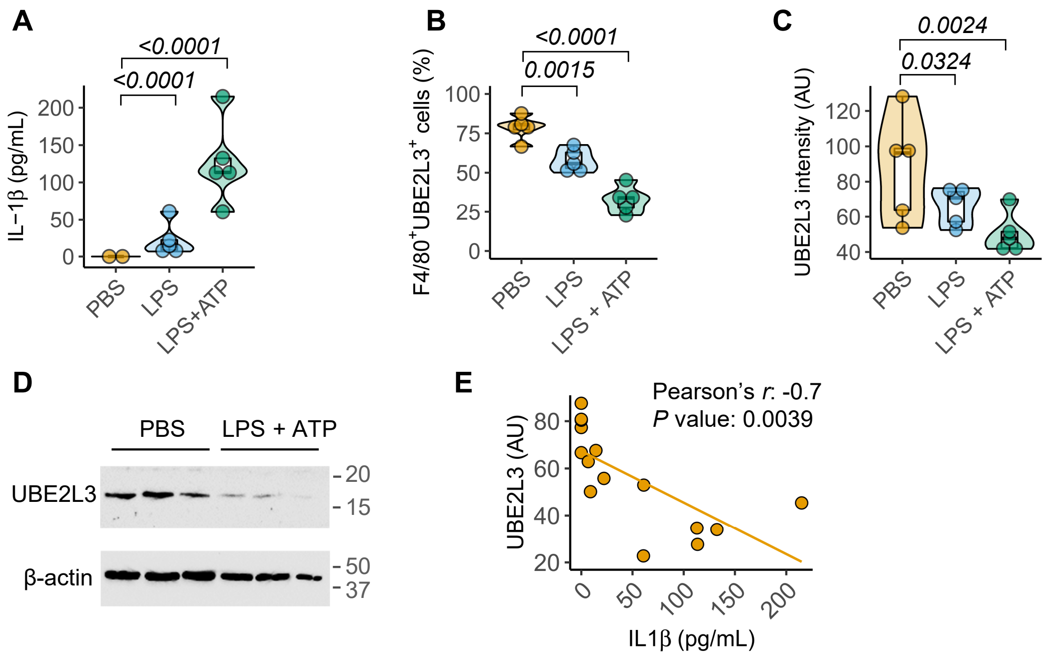


**Figure S1. UBE2L3 is depleted *in vivo* upon inflammasome activation. (A)** IL-1β in peritoneal lavage from mice given PBS, LPS (5 µg, 3 h) or LPS+ATP (5 µg 3 h + 50 µmol, 10 min) intraperitoneally. **(B-C)** Flow cytometry-based quantification using antibodies against F4/80 (macrophage marker) and UBE2L3 in peritoneal cells from mice given PBS, LPS or LPS+ATP as in **(A)**. Graphs show the percentage of double-positive cells **(B)** and mean fluorescence intensity in arbitrary units (AU) of UBE2L3 in F4/80+ve cells **(C)**. **(D-)** Representative immunoblots for UBE2L3 in lysates from peritoneal macrophages from mice given PBS or LPS+ATP (5 µg for 3 h + 50 µmol 10 min) intraperitoneally as labelled. Each lane represents a mouse. Data from one of two similar experiments. **(E)** Plot showing negative correlation between IL-1β measured in peritoneal lavage fluid and UBE2L3 staining by flow cytometry (arbitrary units, AU) in peritoneal macrophages from the same mouse. Each dot represents a mouse given PBS, LPS or LPS+ATP as in **A-C**. Each lane in **D**, and dot in graphs represents an individual mouse. Data distribution is depicted with violin, box (25th to 75th percentile, line at median), and whiskers (± 1.5 x IQR). Two-tailed *P* value for indicated comparison from mixed model ANOVA. ns – not significant ($P > 0.05$).

**Figure S2:** *Mishra, Crespo-Puig et al*

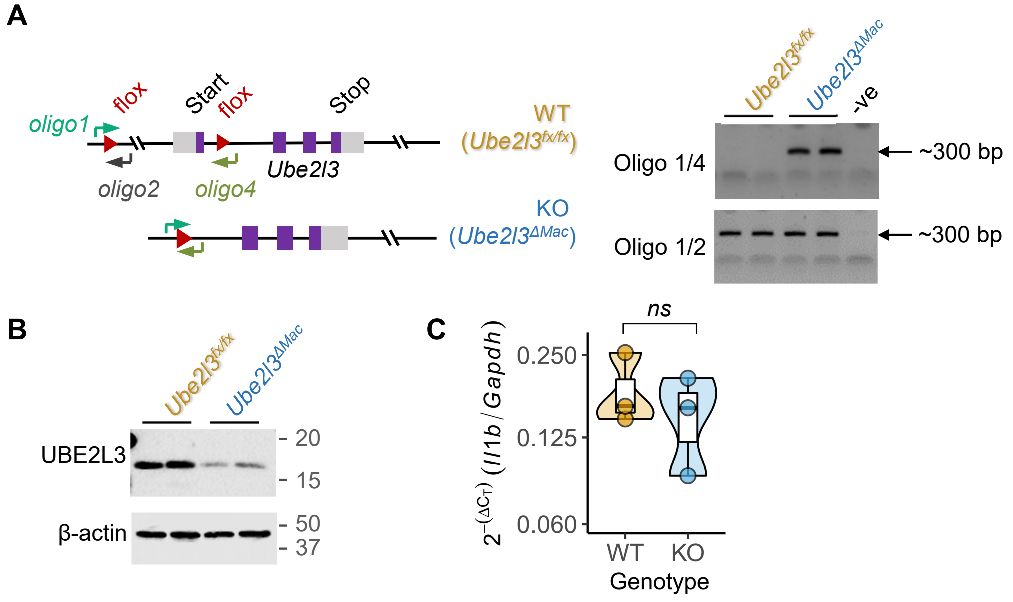


**Figure S2. Generation of *Ube2l3*$^{\Delta Mac}$ mice. (A)** Schematic depiction (left) and PCR-based screening (right) for genotyping WT (*Ube2l3*$^{fx/fx}$) and KO (*Ube2l3*$^{\Delta Mac}$) primary BMDMs given 4-hydroxytamoxifen (2 μM) for 48 h. Oligonucleotide (oligo) primer binding sites are indicated. Oligo 2 binding site is lost after Cre-mediated deletion of exon 1. Oligos 1 and 4 only generate a product under conditions of PCR if homologous recombination has occurred. **(B)** Representative immunoblots from peritoneal macrophages isolated from mice of the indicated genotypes given tamoxifen (80 mg.mL$^{-1}$) orally on three consecutive days. **(C)** Relative expression of *Il1b* normalised to *Gapdh* in peritoneal macrophages isolated from WT (*Ube2l3*$^{fx/fx}$) and KO (*Ube2l3*$^{\Delta Mac}$) mice given tamoxifen. Cells were treated with LPS (250 ng.mL$^{-1}$) for 12 h before preparing RNA for qRT-PCR. Each lane in **B** and dot in **C** represents an individual mouse. Data distribution is depicted with violin, box (25th to 75th percentile, line at median), and whiskers (± 1.5 x IQR). Two-tailed *P* value for indicated comparison from mixed model ANOVA. ns – not significant ($P > 0.05$).

**Figure S3:** *Mishra, Crespo-Puig et al*

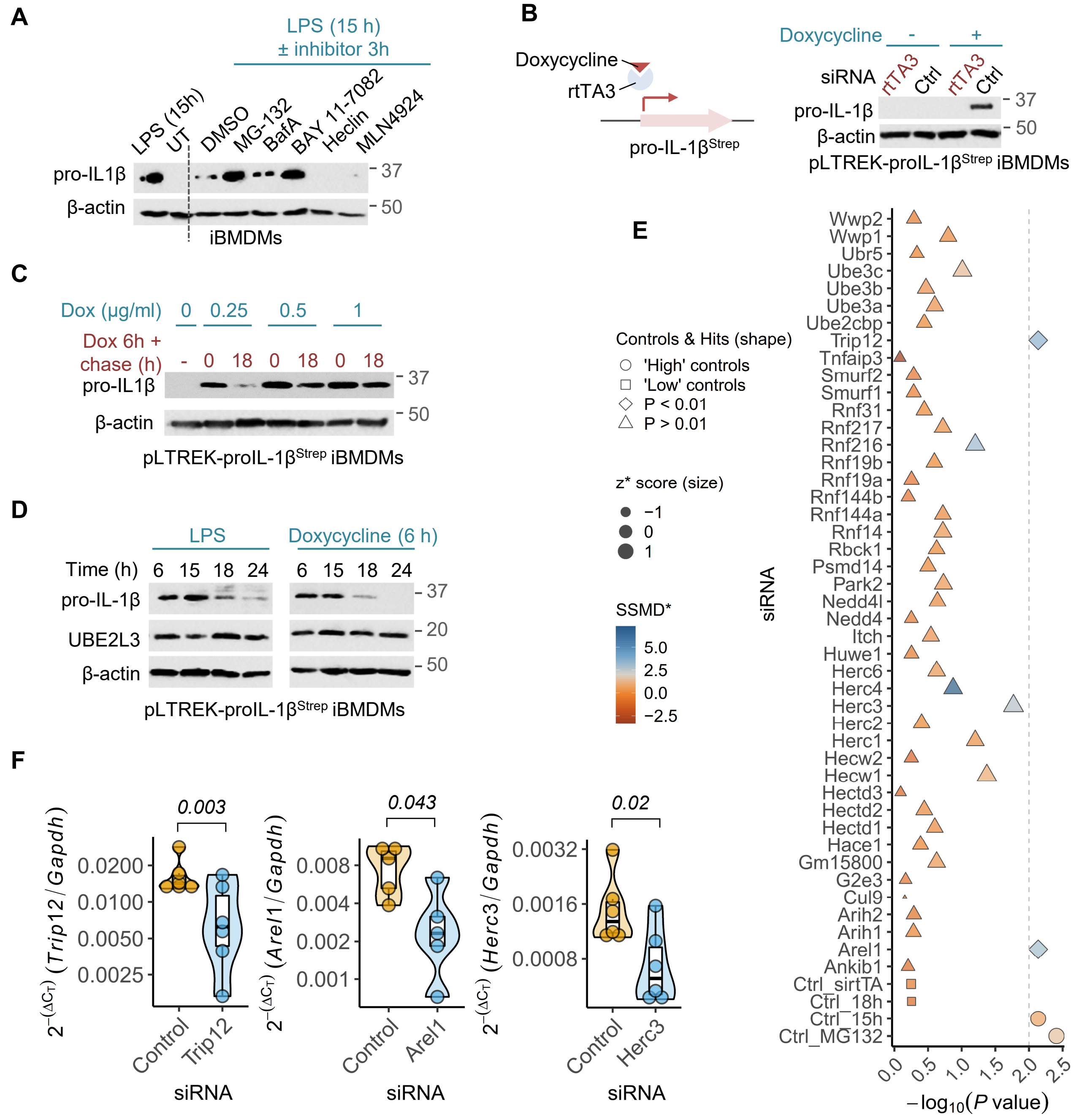


**Figure S3: Validating the siRNA screening approach to identify E3 ligases involved in pro-IL-1β clearance. (A)** Representative immunoblots showing the effect of the indicated inhibitors on the abundance of LPS-induced pro-IL-1β in iBMDMs. Cells were treated with LPS (250 ng.mL$^{-1}$) for 15 h or left untreated (UT) as controls (first two lanes, respectively). Other samples are from cells treated with LPS (250 ng.mL$^{-1}$) for a total of 18 h where the last 3 h included the indicated inhibitors or the solvent DMSO. MG132 (proteasome inhibitor) and BAY 11-7082 (LUBAC and UBE2L3 inhibitor) treatments increased pro-IL-1β abundance as compared to DMSO or other treatments. **(B)** (Left) Schematic showing the transcription factor rtTA3 inducing the expression of pro-IL-1βStrep in the presence of doxycycline in cells stably expressing the pLTREK-proIL-1βStrep plasmid. (Right) Representative immunoblots from lysates of pLTREK-proIL-1βStrep iBMDMs transfected with non-targeting control (Ctrl) or rtTA3 siRNA for 72 h, and then treated with doxycycline for 6 h. **(C)** Representative immunoblots from pLTREK-proIL-1βStrep iBMDMs treated with the indicated concentrations of doxycycline for 6 h, washed and incubated for either 0 or 18 h as labelled. **(D)** Representative immunoblots from pLTREK-proIL-1βStrep iBMDMs treated with LPS (250 ng.mL$^{-1}$) or doxycycline (500 ng.mL-1) as indicated, followed by chase for the indicated times. (legend continues on next page)

**Figure S3:** *Mishra, Crespo-Puig et al (cont.)*

**Figure S3: Validating the siRNA screening approach to identify E3 ligases involved in pro-IL-1β clearance. (cont.)** **(E)** Results from siRNA screen to identify HECT and RBR E3 ligases involved in pro-IL-1β turnover. Size of the symbols indicate mean of z* scores from three independent repeats. FDR-adjusted *P* values for comparison for each of the siRNA with the 'Ctrl_18h' group are shown by shapes of symbols ($P < 0.01$ or $P > 0.01$); 'high' and 'low' controls are also indicated by shapes of symbols. The divergent colour scheme indicates SSMD* scores and the dotted horizontal line indicate $P = 0.01$ cut-off on a -log10() scale. Trip12 and Arel1 scored above thresholds. **(F)** qRT-PCR showing the efficacy of silencing of *Arel1*, *Trip12* and *Herc3* transcripts. iBMDMs were transfected for 72 h with non-targeting control or siRNA against AREL1, TRIP12 or HERC3 as labelled. *Gapdh* was used as normalising control. Images in **A-D** represent experiments performed at least three times. In **F** each dot represents a biologically independent experiment. Data distribution is depicted with violin, box (25th to 75th percentile, line at median), and whiskers (± 1.5 x IQR). P values for indicated comparisons from mixed effects ANOVA.

**Figure S4:** *Mishra, Crespo-Puig et al*

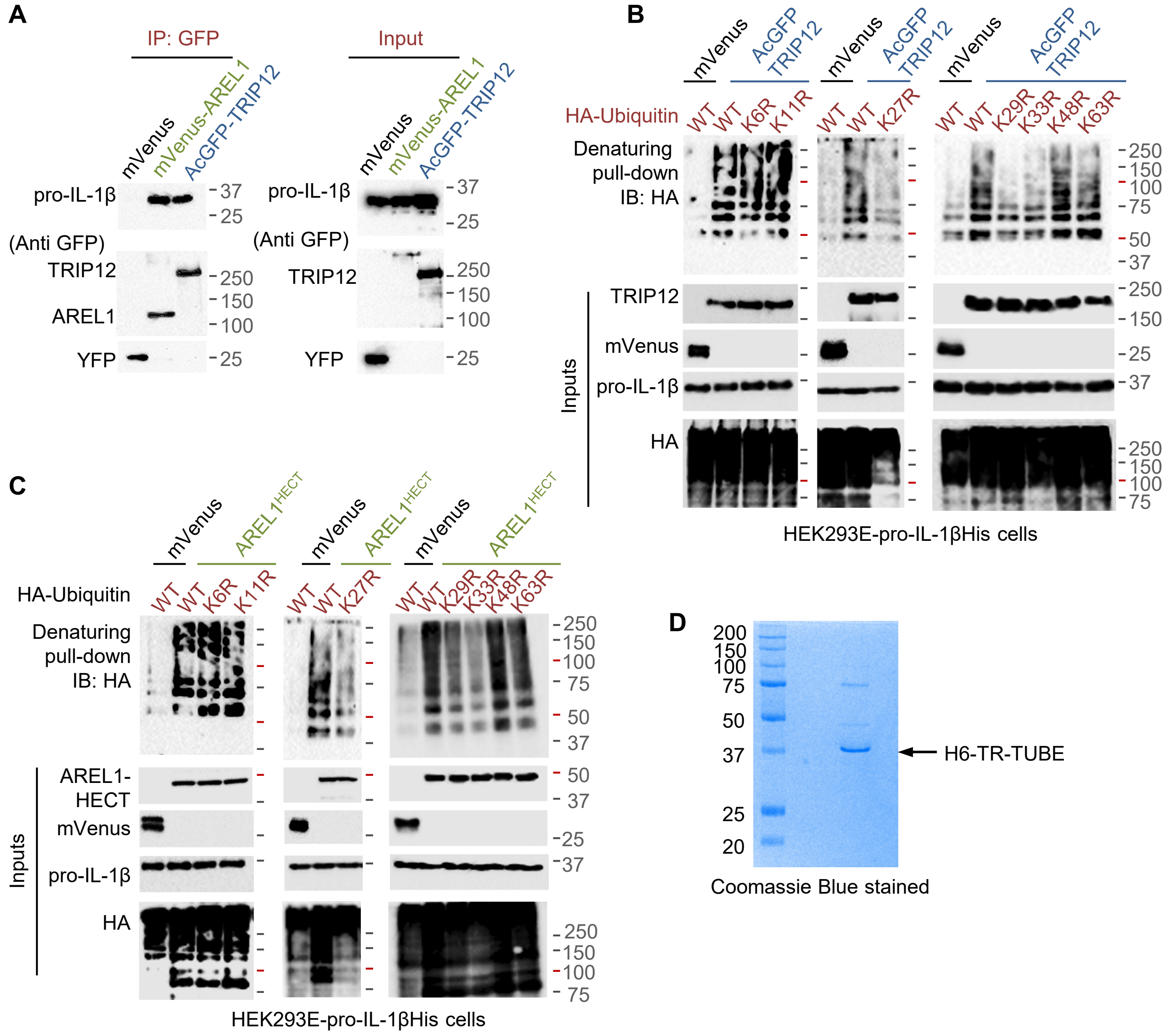


**Figure S4. AREL1 and TRIP12 ubiquitylate pro-IL-1β in the 'pro' domain. (A)** Representative images of immunoprecipitation (IP) and immunoblot (IB) experiments to assess the interaction between pro-IL-1β and TRIP12 or AREL1. HEK293E cells were transfected with plasmids encoding mVenus-AREL1$^{Myc}$ or AcGFP-TRIP12 or mVenus as negative control followed by IP with anti-GFP antibody and IB with pro-IL-1β or GFP antibodies as labelled. Expression of proteins is shown on right (Input). **(B-C)** Representative immunoblots from Ni-NTA pull-downs of pro-IL-1βHis under denaturing conditions (8 M urea-containing buffers) to assess the type of ubiquitin chains covalently added on pro-IL-1β by TRIP12 **(B)** or AREL1 **(C)**. HEK293E cells stably expressing pro-IL-1βHis were transfected with HA-tagged wildtype or the indicated K→R mutants of ubiquitin along with AcGFP-TRIP12 **(B)** or AREL-HECT$^{Myc}$ **(C)** or mVenus as negative control. Expression of proteins is shown below (Inputs). **(D)** SDS-PAGE of purified recombinant hexahistidine-tagged TR-TUBE (~2 μg) followed by Coomassie staining. Images in **A-C** from experiments performed at least three times.

**Summary & Model:** *Mishra, Crespo-Puig et al*

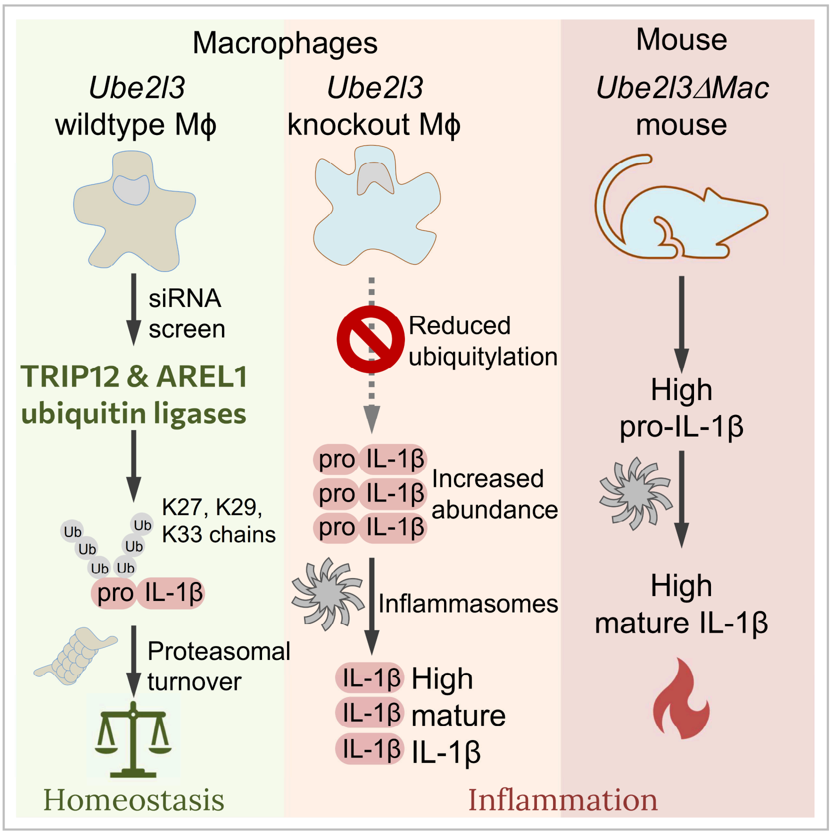


- Conditional deletion of *Ube2l3* ubiquitin conjugating enzyme *in vivo*
- Reduced pro-IL-1β turnover in *Ube2l3*$^{\Delta Mac}$ macrophages and mice
- Elevated IL-1β release and inflammation in *Ube2l3*$^{\Delta Mac}$ mice
- TRIP12 and AREL1 ubiquitin ligases destabilise pro-IL-1β
